# Chd4 remodels chromatin to control retinal cell type specification and lineage termination

**DOI:** 10.1101/2025.01.09.632287

**Authors:** Sujay Shah, José Alex Lourenço Fernandes, Suma Medisetti, Pierre Mattar

**Affiliations:** Regenerative Medicine Program, Ottawa Hospital Research Institute (OHRI), Ottawa, ON, K1H 8L6; Department of Cellular & Molecular Medicine, University of Ottawa, Ottawa, ON, K1H 8M5

**Keywords:** Nucleosome remodelling, Neural progenitor, Retinal Development, Neurogenesis, Temporal development, NuRD complex, Gliogenesis, Photoreceptors

## Abstract

During development, neural progenitor cells modify their output over time to produce different types of neurons and glia in chronological sequences. Previous studies have shown that epigenetic processes play a crucial role in regulating neural progenitor potential, but the underlying mechanisms are not well understood. Here, we hypothesized that nucleosome remodelling would regulate the competence transitions of retinal progenitors. We generated retina-specific conditional knockouts (cKOs) in the key nucleosome remodelling enzyme *Chd4*. *Chd4* cKOs overproduced early-born retinal ganglion and amacrine cells. Postnatally, later-born rod photoreceptors were drastically underproduced. Concomitantly, progenitors failed to be exhausted at late phases of development and ultimately overproduced Müller glia. To determine how Chd4 regulates the genome, we used cut&run-seq to reveal Chd4 genome occupancy, and ATAC-seq experiments to visualize nucleosome remodelling. These data revealed that genome accessibility was significantly increased at ∼10,000 regulatory elements and ∼4,000 genes in the *Chd4* cKO. Together, these results suggest that Chd4 restricts the genome to repress progenitor identity and promote rod photoreceptor production. Accordingly, multiplexed single-cell transcriptomics demonstrated that deletion of *Chd4* led to markedly divergent gene expression profiles. However, despite overproduction of early fates and underproduction of later-born rods, the perinatal transition between early and late progenitor competence was not altered as determined by birthdating experiments and transcriptomic signatures. Taken together, our data suggest that Chd4-dependent chromatin remodelling regulates cell fate specification, and is also required to terminate retinal neurogenesis, but that it does not regulate the progenitor competence windows that restrict early- vs. late-cell-type production.

**Key findings:**

1) *Chd4* cKOs exhibit a strong shift in neurogenesis, with early-born neurons overproduced and late-born neurons underproduced.
2) We present an epigenetic atlas that combines the occupancy of NuRD proteins such as Chd4 and Mbd3, with nucleosome remodelling data and transcriptomic correlates.
3) We show that NuRD regulates the competence transition that terminates the retinal lineage but not earlier competence transitions, showing for the first time that the epigenetic mechanisms governing retinal competence transitions may vary.

## Introduction

The assembly of neural circuits depends on the choreographed production of a constellation of different types of neurons and glia during development. Neural progenitors generate this cell type diversity in precise numbers and proportions on a tightly regulated developmental schedule. In virtually every lineage, progenitors progressively alter their output over developmental time. For example, most CNS progenitors initially make neurons, but then irreversibly switch to producing glia [1]. Moreover, highly complex sequences of neurons and glia are produced in many regions, including the developing retina and mammalian neocortex. Landmark studies in vertebrate and invertebrate systems have demonstrated that the ordered production of different neurons and glia depends in part upon temporal transitions in progenitor competence states [2].

The vertebrate retina is a classic experimental model with which to understand the cell intrinsic mechanisms that regulate the developmental potential of neural progenitors. The mature retina is composed of 6 different types of neurons and one type of glia. During development, this diversity is generated by multipotent retinal progenitor cells (RPCs) [3–5] that exhibit distinct phases of competence. In each phase, RPCs have the potential to generate multiple cell types simultaneously. RPCs initially produce ganglion cells, horizontals, cones, and amacrine cells. As the generation of these cell-types peaks, rod photoreceptors begin to be produced. In rodents, RPCs undergo a competence transition perinatally. Rods are generated in peak numbers during early postnatal stages, and RPCs also continue to generate amacrines but completely lose their capacity to produce the other early-born cell types. At the end of development, RPCs produce rods, bipolar cells, and finally Müller glia [6–10]. While a single pool of RPCs generates this cell type diversity, RPCs also give rise to ‘neurogenic’ cells that have more restricted proliferative and developmental potential. Retinal neurons are often thought to arise from specific neurogenic precursors, whereas Müller glia are thought to differentiate directly from RPCs without passing through a neurogenic precursor state [11, 12]. This ordered production of retinal cell types is conserved across all vertebrates.

While we know a great deal about the developmental mechanisms that program retinal cell type identities, we know much less about how the competence of retinal progenitor cells is controlled. Cell-extrinsic signals have been shown to regulate progenitor output, although these factors are mainly thought to act as *post-hoc* feedback systems that refine cell type production [13, 14]. Previous work has also identified candidate cell-intrinsic competence determinants, which include transcription factors and microRNAs [8, 15–22]. In both the retina and in other CNS regions, competence transitions have also been shown to depend on heterochromatic determinants, including DNA methylation and the polycomb repressor complex [23–25]. These observations suggest a model where heterochromatic processes might act downstream of transcription factors and microRNAs in order to reinforce their activities. The “decommissioning” of genes associated with early competence could explain the loss of developmental potential (progressive restriction) observed during retinal development. However, it remains unclear how the recruitment of heterochromatic determinants such as polycomb might be regulated during development.

The Nucleosome Remodeling and Deacetylase (NuRD) complex is a strong candidate for integrating dynamically expressed temporal transcription factors with heterochromatic effectors. NuRD has dual enzymatic activities –histone deacetylation and nucleosome remodeling. In neural progenitors, the nucleosome remodeling activity is mainly provided by Chd4, a chromodomain helicase protein [26]. Previous work in the cerebellum has shown that Chd4 is required to decommission genes and regulate higher-order genome looping [27, 28]. In the neocortex, precocious gliogenesis was also observed when the NuRD subunit *Mbd3* was mutated [29], suggesting that NuRD might regulate the transition from neurogenic to gliogenic competence. Moreover, *Chd4* cKOs exhibited a loss in late-born upper-layer cortical neurons, suggesting that Chd4 might additionally regulate earlier competence transitions [26, 30]. Interestingly, the NuRD complex has been shown to interact with several temporal transcription factors linked to progenitor competence including Ikzf1, Casz1, Foxp1, and Lhx2 [31–36]. However, the genetic requirement for Chd4 has not previously been examined in the context of retinal development.

Here we used conditional genetics to study how Chd4 regulates retinal development. We report that Chd4 deficiency has a strong effect on the production of early vs. late -born neurons. *Chd4* conditional knockouts (cKOs) markedly overproduce early born RGCs, whereas late-born rod photoreceptors are underproduced. The failure to generate photoreceptors correlated with RPC accumulation beyond the normal developmental window, leading ultimately to an overproduction of Müller glia. Surprisingly, despite the fact that neurogenesis was skewed towards early-born fates and away from later-born rods, we show that early RPC competence is not prolonged in the Chd4 cKO. Thus, while RPCs depend upon Chd4 and nucleosome remodelling for the temporal transition that terminates the RPC lineage at the end of retinal development, they do not require Chd4 for earlier temporal transitions. Taken together, these data hint that sequential competence transitions are regulated by different epigenetic mechanisms.

## Results

### Chd4 expression during retinal development

We first utilized immunohistochemistry to examine the spatiotemporal expression profile of Chd4 during retinal development. We found that Chd4 was ubiquitously expressed from embryonic day (E) 11.5 through to adult stages (Fig. 1). Expression levels appeared to be relatively constant within the nuclei of Ki67+ RPCs from E13.5 through to postnatal day (P)2 (Fig. 1D-K). However, Chd4 levels became somewhat elevated in postmitotic neurons within the ganglion cell layer (GCL) (Fig. 1D-F). At P0 and P2, elevated expression was also apparent within postmitotic Ki67-negative cells within the outer neuroblastic layer (ONBL) (Fig. 1J-M). In the adult retina, Chd4 expression levels remained particularly elevated within inner nuclear layer (INL) and GCL neurons. Weak expression was also apparent in rod photoreceptors (Fig. 1L-O). To determine how other group II subfamily Chd paralogs are expressed during retinal development, we examined previously published retinal scRNA-seq data [8]. While *Chd4* was expressed at relatively high levels in RPCs, both *Chd3/5* were restricted to postmitotic neurons with negligible expression levels in RPCs (Fig S1). These results indicate that Chd4 is the main group II paralog in RPCs, but that its expression is not temporally dynamic.

**Fig. 1.**
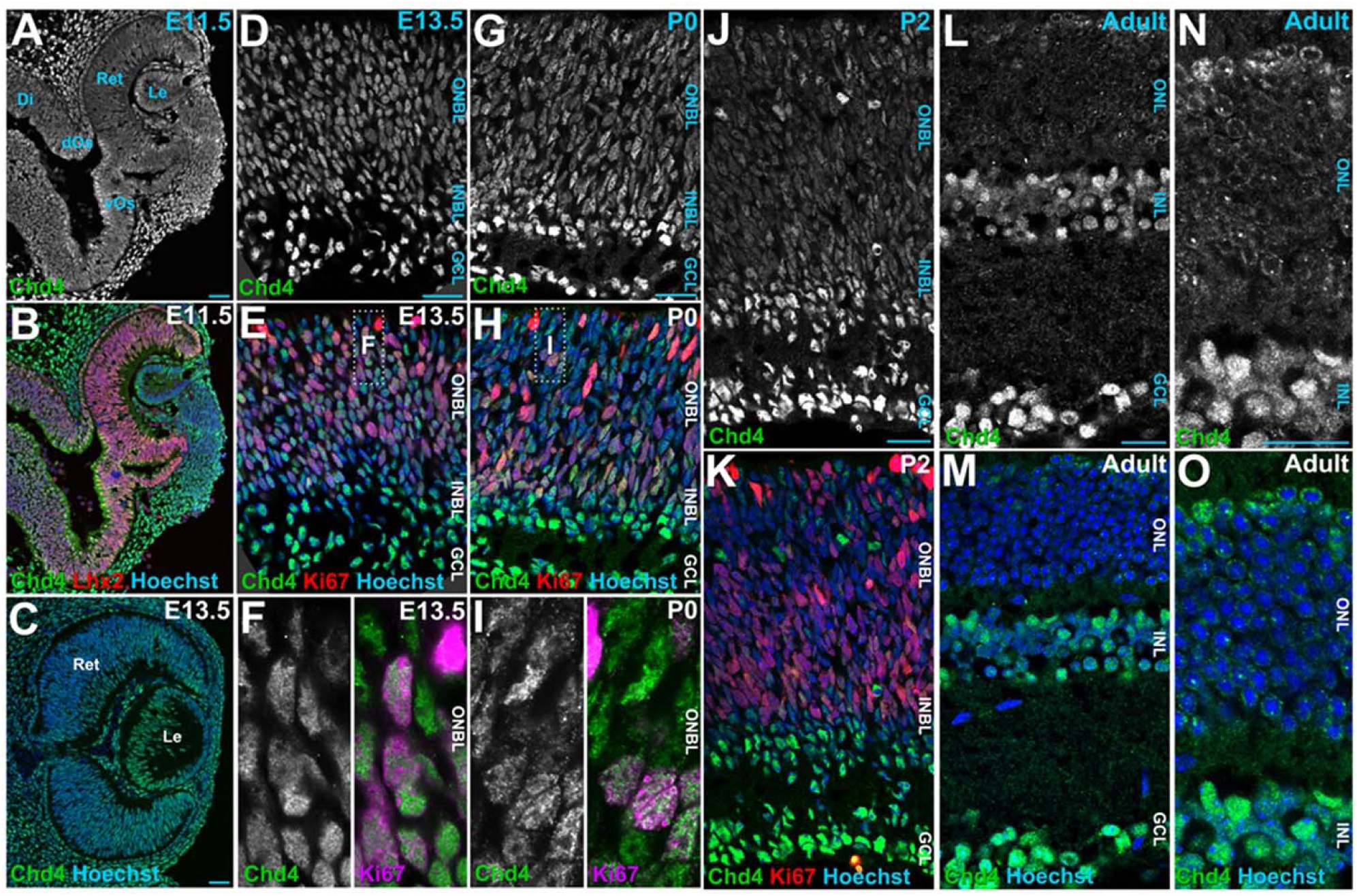
Chd4 expression dynamics during retinal development. (A-B) Immunohistochemistry of wild-type E11.5 retinas with Chd4 (top) and co-staining of Chd4 and Lhx2 (bottom). (C-F) Co-stainings of E13.5 wild-type retinas with Chd4 and Ki67. (G-I) Co-staining of Chd4 and Ki67 in wild-type P0 retinal sections. (J-K) P2 wild-type retinal sections co-stained with Chd4 and Ki67. (L-O). Adult retinal sections stained with Chd4 and Hoechst. Ret: retina; Le: lens; Di: diencephalon; dOs: dorsal optic stalk; vOs: ventral optic stalk; ONBL: outer neuroblastic layer; INBL: inner neuroblastic layer; ONL: outer nuclear layer; INL: inner nuclear layer; GCL: ganglion cell layer. Scale bar = 10 microns.

### Chd4 cKO affects postnatal retinal histogenesis

To determine how chromatin remodelling regulated retinal development, we generated *Chd4* conditional knockouts (cKOs). Since *Chd4* is an essential gene [37], we utilized an allele with loxP sites flanking exons 12-21, which together encode the ATPase/helicase domain [38]. This cassette was deleted using the *Chx10-Cre-EYFP* driver [39], which expresses a Cre-YFP fusion protein in RPCs beginning at ∼E10.5. Towards the end of development, Cre-YFP expression is maintained in bipolars and a few Müller glia. To assess the robustness of this approach, we analyzed Chd4 protein levels in P0 control and *Chd4* cKO retinas. Chd4 protein was efficiently abrogated in cKO retinas (Figure 2A-C).

**Fig. 2.**
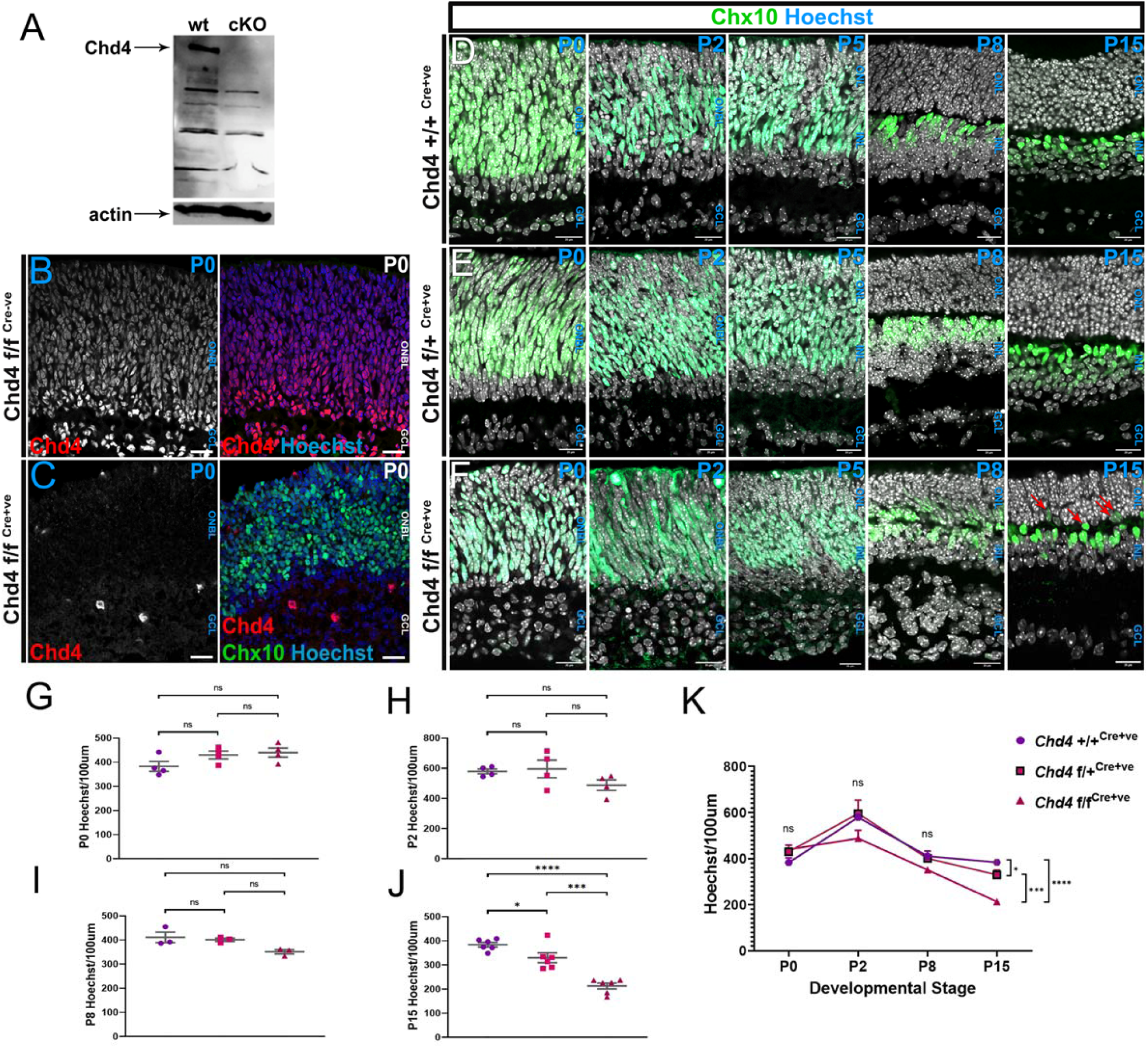
Chd4 is required for retinal histogenesis. (A) Western analysis of Chd4 protein expression. (B-C) Chd4 immunostaining at P1 on wild-type (B) or mutant (mut, *Chd4* f/f ^Cre+^; C). (D-F) Wild-type (wt, *Chd4* +/+^Cre+^), heterozygous (het, *Chd4* f/+^Cre+^) and mutant retinas from P0, 2, 5, 8 and 15 were stained with Hoechst and YFP. Red arrows indicate ectopic YFP+ cells in the ONL. (G-K) Quantification of total Hoechst counts at P0, P2, P8 and P15 as indicated. All data are presented as mean ±SEM. *p<0.05, ***p=0.0001, ****p<0.0001, ns=not significant by one-way ANOVA with Tukey’s multiple comparisons test. Scale bar = 10 microns.

Although the *Chx10-Cre-EYFP* driver is prone to mosaicism, in our study, we observed that an average of ∼70% of cells within the ONBL expressed YFP in perinatal Cre+ animals (Fig. S2), which is similar to the overall proportions of RPCs within the layer (eg. see Fig. 4). Along with the strong reduction in Chd4 protein observed in cKOs, these data indicate that Cre mosaicism was relatively minor in our transgenic animals. We therefore did not exclude any animals or samples in our subsequent analyses.

Next, the effect of *Chd4* ablation was assessed at various stages between E16.5 and P15 (Fig. 2D-F, Fig. S3). *Chd4* cKOs exhibited a markedly expanded GCL along with a poorly formed inner plexiform layer. The distinct neuropil dividing the ONBL and GCL in the wild-type (wt) and conditional heterozygote (chet) was missing in the cKOs (Fig. 2D-F, Fig. S3). However, when the total number of cells was quantified between the three genotypes, no significant changes were observed at P0 or P2 (Fig. 2G, H; Fig. S3). This indicates that the expansion of the GCL does not arise as a consequence of significant retinal overgrowth, nor from premature differentiation of the progenitor pool.

At P8, *Chd4* cKOs exhibited disorganized retinal lamination. Along with the expanded GCL, mutant retinas exhibited a thinned ONL as compared to wt or chets, and additionally exhibited ectopic YFP+ nuclei within the ONL (Fig. 2F). Again, when cells were quantified, no significant difference was observed between wt, chet, and cKO retinas (Fig. 2I). These results indicate that the atypical lamination is not a consequence of altered cell numbers. At P15 the loss of *Chd4* resulted in a hyperplastic ONL containing ectopic YFP+ cells (Fig. 2F, Fig S4). cKO retinas exhibited an approximately 1.5-fold decrease in cell numbers when compared to wt or chet (Fig. 2J). These data suggest that in the cKO, a wave of cell death occurs between P8 and P15 (Fig. 2K), likely corresponding to the wave of apoptosis previously shown to prune supernumerary bipolars and amacrines at ∼P10 [39–42]. Since the proportions of cells present in each retinal layer were already distorted at P8 prior to this wave of cell death, these data collectively indicate that Chd4 balances cell-type production during retinal development.

### Chd4 regulates retinal cell-type production

To understand how the cell composition of the retina was affected, we examined cell-type specific markers at P15, when development is complete. We found that cKO retinas had an almost 2-fold increase in the proportion of Rbpms+ RGCs as compared to controls (Figure 3A-D). To measure amacrines, we counted Pax6+ cells within the INL. cKO retinas displayed an increase in INL amacrines when compared to wt. Pax6+ positive cells were increased almost 4-fold when the GCL was included in the counts, further reflecting the expansion in RGCs (Figure 3C, D). For cones, we found that cKOs exhibited an approximately 2-fold decrease in the percentage of cone arrestin+ cells when compared to wt and chet retinas (Figure 3E, F).

**Fig. 3.**
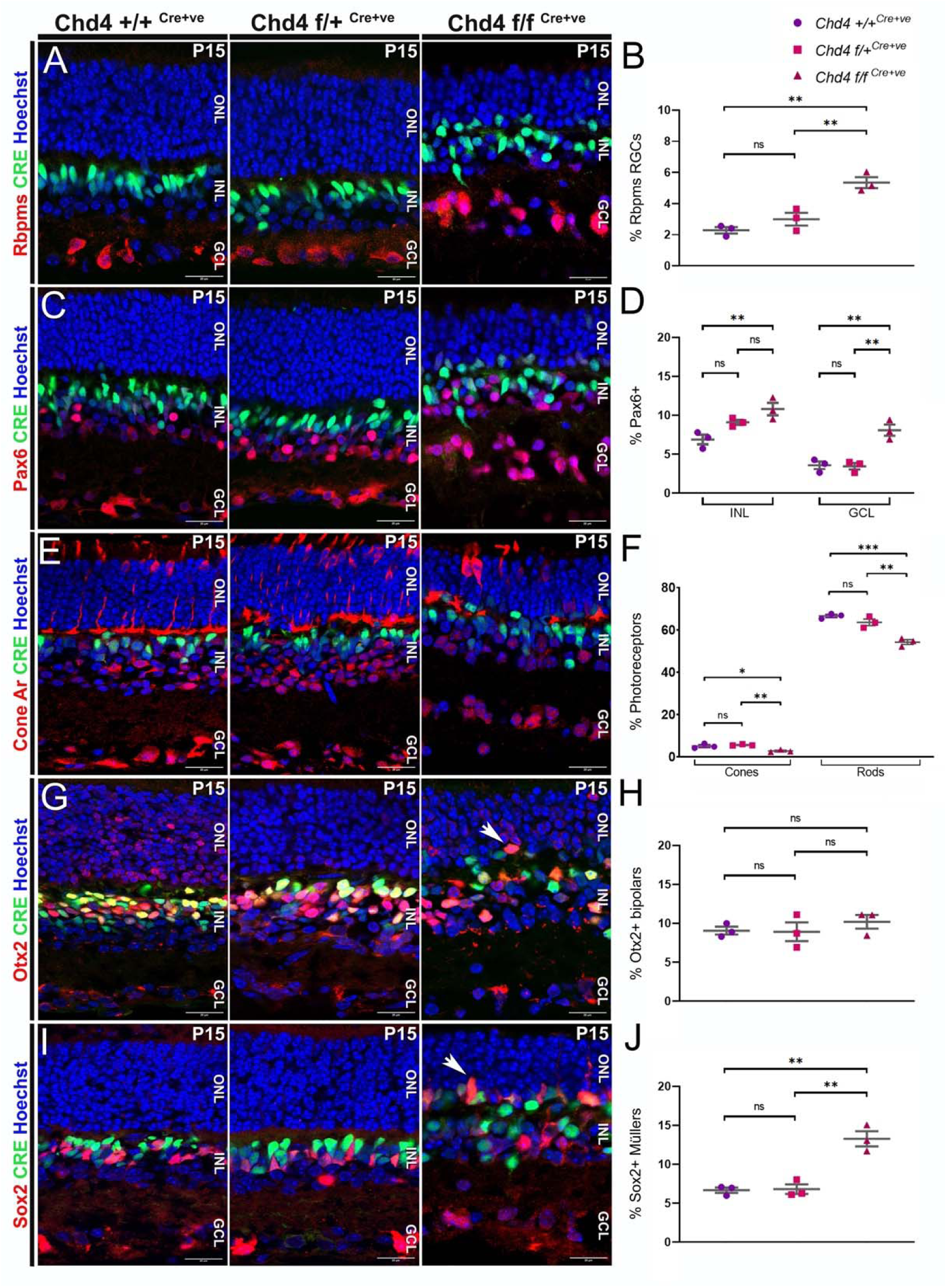
Chd4 regulates retinal cell type composition. (A, B) Rbpms was used to quantify the percentage of RGCs among the three genotypes. (C) Pax6 marks amacrines in the INL and GCL, along with RGCs in the GCL. (D) Quantification of the percentage of Pax6+ cells by layer as indicated. (E) Cone arrestin was used to quantify the percentage of cones among the three genotypes. (F) GFP-negative/cone-arrestin negative ONL cells were counted as rods. (G, H) Brightly positive Otx2 cells were counted as bipolars. Arrows indicate the ectopic presence of Otx2+ bipolars in the ONL. (I, J) Radially polarized Sox2+ cells were counted as Müller glia. Arrows indicate the ectopic presence of Sox2+ glia in ONL. All data are presented as mean ±SEM. p value *p<0.05, **p<0.005, ***p<0.0005, ns=not significant by one-way ANOVA with Tukey’s multiple comparison test. Scale bar = 10 microns.

During postnatal stages, RPCs generate amacrines, rods, bipolars, and Müller glia. In the ONL, we counted marker-negative rods by excluding cone arrestin+ cones and YFP+ bipolars/Müllers. As expected from the thinning of the ONL (Fig. S4), rods were significantly reduced in *Chd4* cKO retinas as compared to wt or chet (Fig. 3E, F). The proportion of strongly Otx2+ bipolar cells did not differ among the three genotypes at P15. However, cKOs exhibited ectopic Otx2+ and YFP+ bipolar cells within the ONL (Figure 3G, H). During retinal development, Müller glia are the latest-born cell-type. Counting radially polarized Sox2+ cells as Müller glia, we observed an almost 2-fold increase in the proportions of glial cells in the cKO, along with their ectopic presence in the ONL (Fig. 3I, J). These results suggest that Chd4 restricts the production of Müller glia to promote the development of rod photoreceptors.

### Chd4-dependent chromatin remodelling does not regulate progenitor proliferation

The observed shifts in cell type composition could potentially be explained by alterations in progenitor proliferation. For example, if self-renewing divisions were undermined, this might lead to overproduction of early-born RGCs, which could prematurely exhaust progenitors - leading to underproduction of late fates. We therefore examined retinas at P0, when RPCs lose the competence to generate early-born cell types, and just prior to the normal peak in rod production [10]. Brn3a staining confirmed that RGCs were increased approximately 2-fold in the *Chd4* cKO (Fig. 4A-D), similar to the increase in RGCs observed at P15. The expansion in early-born neurons was also illustrated via the RGC marker Rbpms, as well as the amacrine marker Tfap2a (Fig. S5). However, we noted that the size of the progenitor pool - as reflected by the expression of the *Chx10-Cre-YFP* transgene – was not altered (Fig. 4A-E).

**Fig. 4.**
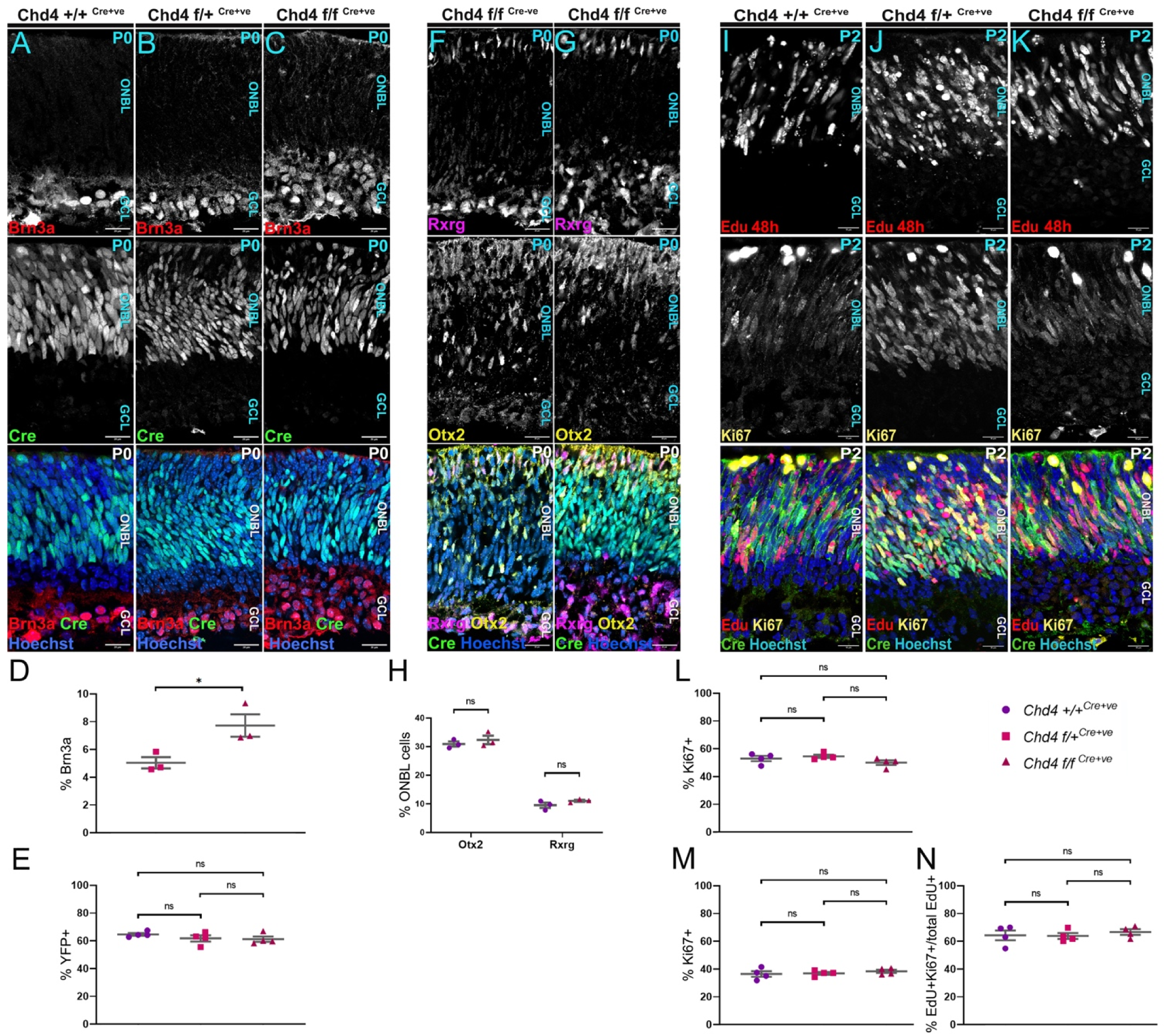
Fate shifts are independent of perinatal alterations in RPC numbers or proliferative properties. (A-C) P0 retinal sections were co-stained with the RGC marker Brn3a and imaged for YFP, which marks RPCs. (D, E) Quantification of the percentage of Brn3a+ cells (D), or YFP+ cells (E). (F-G) P0 retinal sections were co-stained with the cone marker Rxrg, and the photoreceptor precursor marker Otx2. (H) Percentage of Otx2+, and Rxrg+ cells between control and mutant retinas. (I-K) Birthdating was performed by injecting 10mM EdU in P0 pups and harvesting after 48 hrs. (L-N) Percentage of cells that are EdU+ (L), Ki67+ (M), or EdU+Ki67+ as a percentage of total EdU+ cells (N). All data are presented as mean ±SEM. *p<0.05, ns: not significant by one-way ANOVA with Tukey’s multiple comparisons test. Scale bar = 10 microns.

Since these data suggested that RGCs were not overproduced as a consequence of premature progenitor depletion, we hypothesized that RGCs might be produced beyond their normal birth window. Postnatally, only trace levels of RGC production are observed in wild-type mice [10]. We therefore injected EdU at P0. After 2 days, we observed small numbers of newly born EdU+/Brn3b+ RGCs that were often localized apically, suggesting that they were migrating towards the GCL. However, very few of these cells were observed, and there was no marked difference between controls and cKOs (Fig. S6). We conclude that RGC generation is not prolonged at postnatal stages. These data concur with the marked expansion of the GCL already observed by E16.5 and P0 in the *Chd4* cKO (Fig. 2, 4, Fig. S3), as well as scRNA-seq data (see below), which together indicate that supernumerary RGCs are mainly produced during embryogenesis within their normal birth window.

While these data suggest that the progenitor pool was not prematurely depleted, we hypothesized that RGCs might increase at the expense of the rods and cones that are generated during embryonic stages. We therefore counted Otx2+ photoreceptor precursors at P0, as well as Rxrg+ cones. Surprisingly, we found that proportions of early-born photoreceptors in *Chd4* cKO retinas were comparable to controls (Figure 4F-H), suggesting that the decrease in cone arrestin+ cells observed at P15 might reflect subsequent defects in cone differentiation or survival. Thus, while RGCs expand and photoreceptors contract in number, these changes appeared not to be linked to a common fate decision.

Since rod photoreceptor production normally peaks between P0 and P2 [10], we reasoned that RPCs might exhibit defects in the generation of postmitotic daughter cells or altered cell cycle progression that could undermine rod production at these stages. To examine RPC proliferation in more detail, we examined mitotic cells using phospho-histone H3. Again, there was no difference between the genotypes (Fig S2A-D). Next, to assess whether loss of Chd4 influenced progenitor self-renewal, we injected EdU to mark S-phase cells at P0. At P2, RPCs were co-stained for EdU and Ki67, which marks proliferating cells. Both EdU and Ki67 were comparable between the three genotypes (Fig. 4I-M). Double-labelled EdU+/Ki67+ RPCs that had undergone self-renewal were also not significantly different (Fig. 4I-N). These data indicate that *Chd4* cKO did not affect progenitor proliferation dynamics at perinatal stages in agreement with lack of differences in overall cell numbers observed up to P8. However, previous work in the developing neocortex had shown that loss of Chd4 led to elevated cell death [26]. We therefore performed the TUNEL assay at P0. However, we observed no changes in apoptotic frequency in the *Chd4* cKO versus controls at this stage (Fig S2). Thus, while *Chd4* cKOs exhibited distorted cell-type proportions that could have arisen as a byproduct of premature RPC exhaustion or cell death, such effects were not evident perinatally during the normal peak of rod production.

### Loss of Chd4 results in divergent transcriptomic profiles

To further examine how *Chd4* mutation shifts the transcriptional state of RPCs, we performed scRNA-seq at P1, by which point RPCs have normally undergone the transition between early and late competence. To avoid batch effects, we used the Multi-seq barcoding approach [43], allowing us to compare biological replicates for 3 *Chd4* cKOs versus 3 littermate control retinas processed together within the same 10X Genomics Chromium well. Cells were sequenced to a depth of 23,190 reads per cell and a median of 1,980 genes per cell for an estimated sequencing saturation of 49.6%. After demultiplexing and removing low-quality cells and doublets, our dataset retained 9,776 cells, with 2,152 control cells and 7,624 cKO cells. Next, cell-types were annotated using scDeepSort [44] to perform unsupervised label-transfer based on a previously published retinal scRNA-seq atlas [8].

Next, we visualized the data using Uniform Manifold Approximation and Projection (UMAP; Fig 5A). We noted that cells annotated as ‘late RPCs’ formed a wheel-like structure, from which a neurogenic ‘stem’ emerged, followed by a bifurcation towards amacrine cells, or alternatively towards photoreceptor precursors. Marker gene expression confirmed the fidelity of the cell-type annotation (Fig. S7). Next, we examined each replicate (pup) individually (Fig. 5B). To confirm the genotype of each barcode, we examined how *Chd4* and its paralogs were expressed in each replicate. Focusing on RPCs, we observed that *Chd4* cKO cells exhibited a significant reduction in *Chd4* transcription as compared to controls, but that *Chd4* was not eliminated (Fig. 5C). However, this was expected, since the loxP-flanked cassette does not excise the 3′ end of the gene [38]. Since *Chd4* cKO has been shown to lead to a compensatory upregulation in its paralogs *Chd3* and *Chd5* [27, 45], we additionally examined these transcripts and found that both were significantly upregulated in *Chd4* cKO samples as expected (Fig. 5C). Immunohistochemistry confirmed that Chd3 protein upregulated in *Chd4* cKO RPCs, further validating these observations (Fig. S8).

**Fig. 5.**
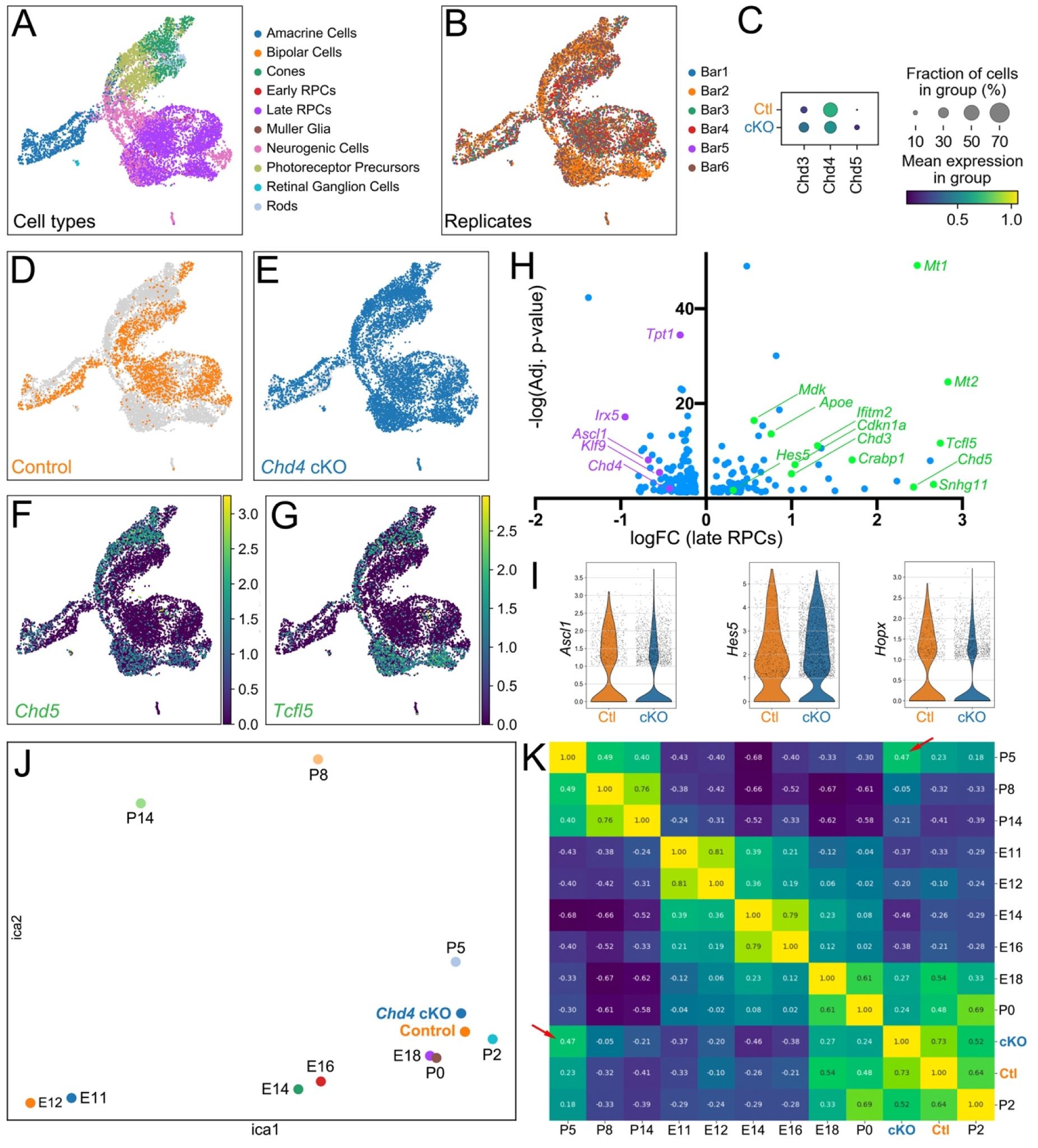
Loss of *Chd4* leads to global transcriptional dysregulation and a slight temporal acceleration. (A) Leiden UMAP clustering of 18,815 single cells from P1 control (n=3) or cKO (n=3) littermates. Cell types were annotated via unsupervised label transfer according to marker genes published in a previous study [8]. (B) UMAP projection of each demultiplexed sample. (C) Dotplot comparing the expression of *Chd4* and its paralogs in late RPCs from control versus cKOs. (D-E) Comparison of control versus cKO cells. (F-G) UMAP projection of the expression of *Chd5* (F) and *Tcfl5* (G). (H) Volcano plot of differentially expressed genes (DEGs) from late RPCs in *Chd4* cKO samples vs control (adj. p-value <0.05; LogFC > 0.4). (I) Violin plots comparing *Ascl1*, *Hes5*, and *Hopx* expression in wt vs. cKO late RPCs. (J) Independent Component Analysis (ICA) comparing wt and cKO scRNA-seq data with a published retinal RNA-seq atlas [8]. (K) Pairwise comparison correlation matrix heatmap of the ICA analysis. Arrows indicate the elevated correlation between the cKO dataset and the P5 samples of the retinal atlas.

Next, we visualized *Chd4* cKOs versus controls. Control cells from each replicate clustered together in UMAP space (Fig. 5D). *Chd4* cKO cells overlapped with control cells, but were additionally shifted into novel parallel clusters that did not contain control cells (Fig. 5E). We found that significantly upregulated genes - including *Chd5* (Fig. 5F), and *Tcfl5* (Fig. 5G), were expressed only in the novel clusters that appeared in *Chd4* cKO samples, but not in clusters occupied by control cells. These data likely indicate that our *Chd4* cKO samples exhibit marked alterations in gene expression across the full developmental trajectory, and that these changes are observed despite probable mosaicism in one of the *Chd4* cKO samples, as well as compensation from *Chd3* and *Chd5* paralogs.

We next identified differentially expressed genes (DEGs; Supplemental datafile 1). Focusing specifically on RPCs, *Chd4* cKOs exhibited both downregulated and upregulated DEGs (Fig. 5H). Downregulated DEGs included the transcription factor *Irx5*, which is involved in the specification of bipolar cell subtypes [46] and the proneural gene *Ascl1* which is necessary for rod and bipolar cell production [47] (Fig. 5I). In accordance with the overproduction of Müllers, upregulated genes included genes such as *Apoe*, *Cdkn1a*, *Hes5*, *Mt1*, *Mt2*, which are glial markers or determinants (Fig. 5H, I). Other upregulated genes included *Ifitm2*, *Tcfl5* and *Snhg11*, all of which were all previously observed to upregulate in the *Chd4* cKO neocortex [45].

Next, we focused on using the scRNA-seq data to address whether there might be a temporal shift in the absence of *Chd4* -dependent chromatin remodelling. Although *Chd4* cKOs exhibit drastic changes in cell composition, major changes in the developmental trajectories of *Chd4* cKOs were not evident. For example, we did not observe the persistence of an RGC neurogenic trajectory. Like other postnatal scRNA-seq datasets [8, 48], we captured very few RGCs in our experiment, albeit that almost all of the recovered RGCs came from the *Chd4* cKO samples. Similarly, while *Chd4* cKOs overproduce Müllers and glial genes were upregulated, unsupervised annotation did not reveal Müller glia already present in the P1 dataset. As Müller production normally peaks between P2 and P5 [10], these data indicate that Müllers do not differentiate prematurely.

Despite lack of obvious temporal shifts in cell type production, we reasoned that a shift in the developmental stage of *Chd4* cKOs could be evaluated by comparing our dataset against other timepoints. We therefore integrated our dataset with an existing developmental scRNA-seq atlas [8]. Using Independent Component Analysis (ICA), we found that most of the timepoints in the published scRNA-seq atlas were arrayed in a logical continuum, with earliest embryonic samples at the origin, perinatal samples differing most along the first component (ICA1), and later postnatal samples differing most along the second component (ICA2; Fig. 5J). In accordance with expectations, both P1 *Chd4* cKO and littermate control samples were localized near to P0 and P2 samples from the retinal atlas. However, we observed that the *Chd4* cKO samples were slightly shifted towards P5 samples, while littermate controls were located closer to P0 and P2 samples. To better visualize this potential shift, we plotted the integrated dataset in a comparison matrix (Fig. 5K). Both P1 control and cKO samples correlated most closely with P2 samples (control: 0.64; *Chd4* cKO: 0.52). Strikingly, *Chd4* cKOs correlated more strongly with P5 samples (0.47) than they did with E18.5 (0.27) or P0 (0.24) samples. By contrast, littermate control samples correlated more strongly with E18.5 (0.54) or P0 (0.48) samples and much less well versus P5 (0.23) as would be expected. Thus, while a shift in temporal identity could not easily be discerned with respect to UMAP trajectories, both global gene expression and DEG signatures suggest that *Chd4* cKOs may be *slightly* accelerated in their temporal state.

### Chd4 regulation of chromatin occupancy and accessibility in RPCs

Next, we wished to determine how Chd4 regulates the genome. To examine the genome occupancy of Chd4, we performed cut&run-seq on P1 wild-type and cKO retinas using a validated Chd4 antibody [27, 45]. We additionally examined Mbd3, which is specific to the NuRD complex. Visual comparison of these datasets revealed correspondence between Chd4 and Mbd3 (Fig. 6A). Across the genome, Chd4 occupied ∼10,000 peaks in wild-type retinas, which was comparable to peak numbers observed in the neocortex and cerebellum [27, 45]. Mbd3 occupied ∼3,500 peaks, with most of these peaks co-occupied by Chd4 (Fig. S9A-C). In *Chd4* cKO retinas, Chd4 and Mbd3 peak numbers were drastically reduced to ∼2,500 and 1,000, respectively (Fig. S9A). When compared to published retinal ChIP-seq data [49], we found that ∼2/3 of the Chd4 peaks localized to gene promoters marked by H3K4me3 (Fig. S9B).

**Fig. 6.**
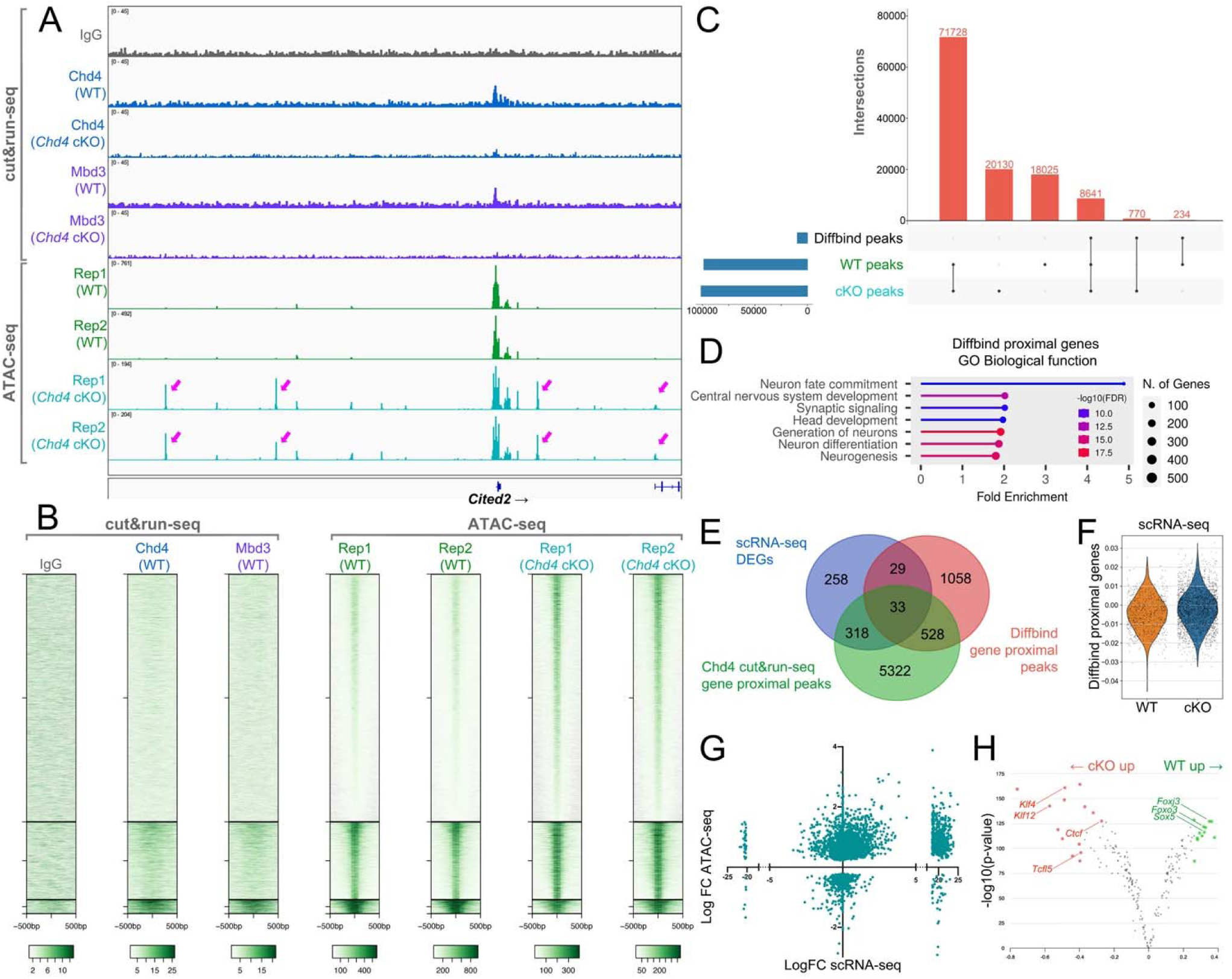
*Chd4* restricts chromatin accessibility in RPCs. (A) Cut&Run-seq and ATAC-seq tracks showing called peaks at *Cited2* locus from P1 control and *Chd4* cKO RPCs. Arrows indicate ectopic peaks present in the cKO samples but not in control. (B) Heat map of the alignment of ATAC-seq datasets of sorted RPCs from control and *Chd4* cKO samples compared against our Cut&Run-seq dataset. Plots are centered on the differentially accessible regions (DARs) of cKO RPCs analyzed by Diffbind. (C) Upset plot comparing the peak intersections between control, cKO, and DARs. (D) GO terms analysis of Diffbind proximal genes. Diffbind proximal peaks were assigned to genes located within 5kb upstream and 1kb downstream of the peak. (E) Integration of DEGs from the scRNA-seq analysis with genes associated with gene-proximal DARs and Chd4 Cut&Run-seq gene-proximal peaks. (F) Violin plot of expression of genes associated with a proximal DAR in our scRNA-seq dataset, comparing control versus *Chd4* cKO RPCs. (G) LIMMA analysis integrating our scRNA-seq with ATAC-seq to compare changes in gene accessibility against fold-change in transcription. (H) Footprinting analysis on accessible peaks using the TOBIAS algorithm.

To directly visualize the nucleosome remodeling activity of Chd4 in RPCs, we performed ATAC-seq on 2 wt and 2 cKO littermates, by sorting YFP+ RPCs marked by the *Chx10-Cre-EYFP* transgene at P1. Inspection of the loxP-flanked cassette revealed almost complete excision in the cKO, validating the sorting strategy (Fig. S10). To identify differentially accessible regions (DARs) in *Chd4* cKO RPCs, we next performed diffbind analysis (Fig 6B) yielding approximately 10,000 DARs between control and mutant RPCs. Most DARs exhibited increased accessibility (Fig. 6A-C). While the NuRD complex has previously been shown to decommission some regulatory elements, we found that most DARs were still nominally accessible in the control datasets (Fig. 6C). More surprisingly, most of these DARs exhibited little Chd4/NuRD complex occupancy (Fig. 6B), indicating a probable “kiss-and-run” transient interaction or indirect regulation. By contrast, DARs that were reduced in accessibility in *Chd4* cKOs appeared to be directly bound, suggesting that loss of the NuRD complex footprint might drive the effect.

To determine how differential accessibility might relate to gene expression, we performed peak-to-gene annotation. We selected only gene-proximal peaks (within 5 kb upstream and 1 kb downstream of the gene body inclusive) in order to filter the overall peak number. Gene proximal DARs were associated with ∼1,600 genes (Supplemental datafile 1). Gene ontology analysis showed that neuron fate commitment, neurogenesis and neuron differentiation were highly enriched (Fig 6D), suggesting potential misregulation of these processes.

To determine how changes in accessibility affect gene expression, we examined genes associated with proximal DARs in our scRNA-seq dataset. We found that DAR-associated genes overlapped with only ∼10% of DEGs. However, more extensive overlap was observed with Chd4 cut&run-seq peaks, with more than half of the DEGs and approximately a third of DAR-associated genes directly occupied by Chd4. Additionally, 33 target genes were common between all three of the datasets (Fig. 6E; Supplemental datafile 1), including genes such as *Cited2* (Fig. 6A), *Mbln2*, *Jund*, and *Plagl1* (Fig. S11). In accordance with these observations, when we generated a gene scoring module for genes associated with a proximal DAR, we found a slight but significant upregulation of this gene score in the *Chd4* cKO (Fig. 6F), suggesting that changes in accessibility correlate with transcription. We next measured accessibility across gene bodies using LIMMA, which allowed us to integrate accessibility data for each gene across all associated peaks rather than on an individual peak-by-peak basis, where the summed effects of multiple peaks can be difficult to infer. Comparing fold-changes in gene accessibility versus fold-changes in transcription, we observed a significant (P<0.0001) positive correlation between increased accessibility and transcriptional upregulation across the genome (Fig. 6G).

Lastly, we performed footprinting analysis on accessible peaks in order to identify differential transcription factor occupancy using the TOBIAS algorithm (Fig. 6H) [50]. Using this approach, we found that Ctcf was one of the overrepresented motifs in *Chd4* cKO RPCs. This indicates that *Chd4* might also be involved in regulating RPC chromatin architecture where the loss of Chd4 might lead to increased recruitment of Ctcf to sites that are typically inaccessible, resulting in disorganization of genome looping as previously shown in cerebellar granule cells [51]. Additionally, Tcfl5 motifs were also overrepresented in mutant RPCs corroborating increased *Tcfl5* transcript levels observed in the scRNA-seq data. Taken together, these data suggest that Chd4 may have a broad role in restricting nucleosome accessibility and consequent transcription across the genome, which might stabilize the RPC identity.

### Chd4 regulates neurogenic competence at late stages

While rod photoreceptors are markedly underproduced in the *Chd4* cKO, our analyses suggest that RPC proliferation and photoreceptor specification proceed normally between embryonic and early postnatal stages. Thus, we wondered whether *Chd4* was required for rod production only during postnatal stages. Since cell cycle parameters are unaffected between P0 and P2, we injected EdU into P1 pups and visualized these birthdated cells at P15. In accordance with the notion that *Chd4* cKOs underproduce rods postnatally, the proportion of EdU+ cells in the ONL was significantly reduced in mutant retinas compared to controls (Figure 7A-C). Conversely, we observed a significant increase in the proportion of EdU+ cells in the INL of the mutant retinas (Figure 7D). These data show that Chd4 biases RPCs to favor rod photoreceptor production over other cell fates. This interpretation was corroborated via retroviral lineage tracing experiments where P0 RPCs appeared to generate more glia at the expense of rod photoreceptors in the *Chd4* cKO without effects on clone size (Fig S12).

**Fig. 7.**
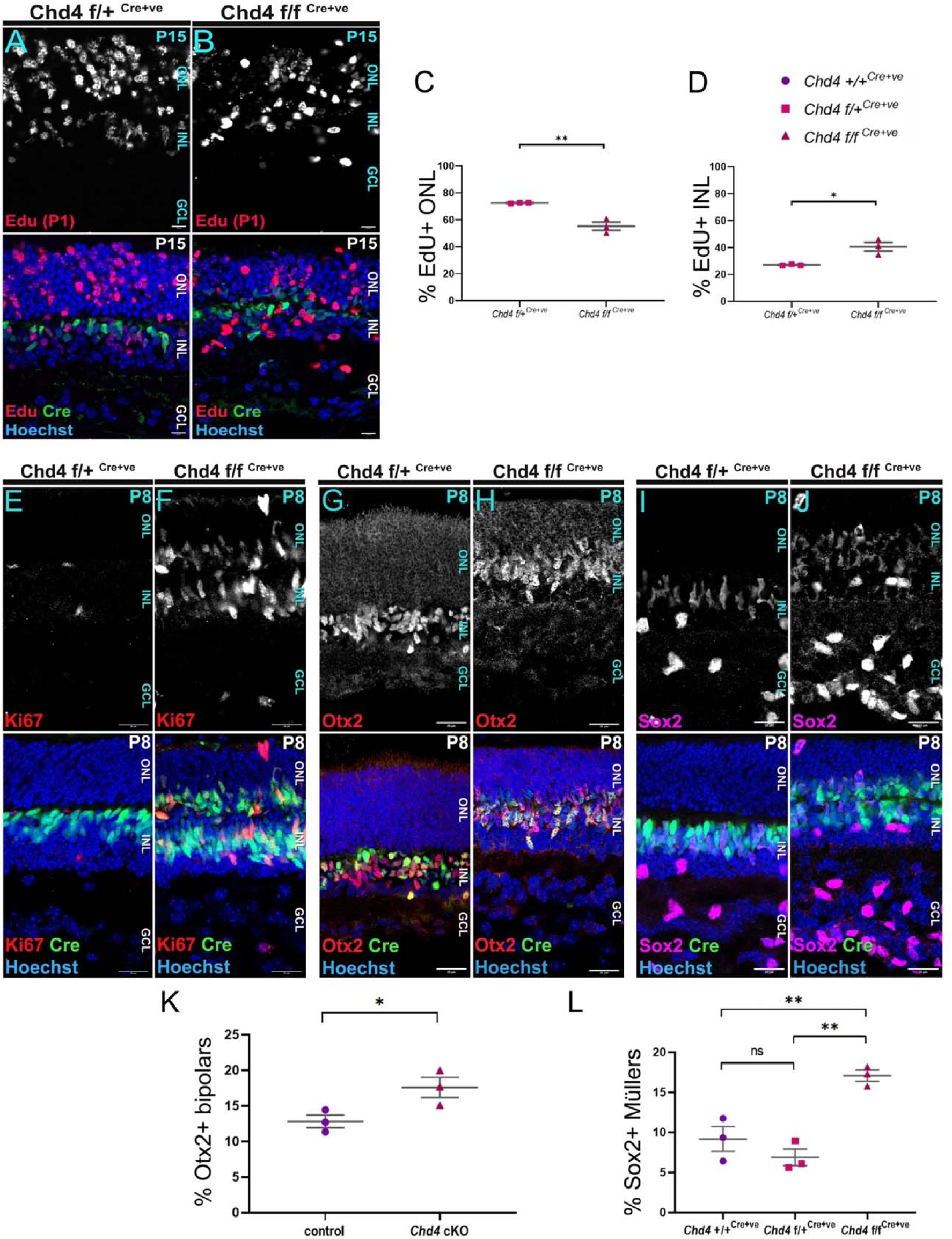
Chd4 is required to terminate the retinal lineage. (A, B) EdU was injected at P1 and retinas were harvested at P15. (C, D) Quantification of the proportion EdU+ cells divided between the INL (C) and ONL (D). (E-J) P8 central retinal sections stained for the proliferation marker Ki67 (E, F), the bipolar marker Otx2 (G, H), or the RPC/glial marker Sox2 (I, J). (K, L) Quantification of the percentage of brightly Otx2+ bipolar cells (K), or radially polarized Sox2+ Müllers (L). All data are presented as mean ±SEM. p value *p<0.05, **p<0.005, ns=not significant by two-tailed unpaired Student’s t test or one-way ANOVA with Tukey’s multiple comparison test. Scale bar = 10 microns.

Since Müller glia arise directly from RPCs without passing through a neurogenic precursor state [11, 12], we next examined retinal development at P8 in order to determine how neurogenesis is affected at late stages. Retinal sections again exhibited the ectopic presence of YFP-positive cells in the ONL in mutant retinas compared to controls. Ki67 staining revealed the persistence of cycling RPCs within the central retina beyond their normal temporal window (Figure 7E, F). To examine bipolars, we stained for Otx2, revealing a significant increase in bipolars at P8 (Fig. 7G, H, K), which probably die back by P15. Similarly, Sox2-positive Müllers were significantly increased in the cKO retinas compared to controls (Figure 7I, J, L). Taken together, the persistence of cycling progenitors along with the marked underproduction of rods suggest that RPCs prematurely lose the competence to generate rod photoreceptors and fail to be exhausted, leading ultimately to the overproduction of alternative fates.

## Discussion

Although temporal transitions play a critical role in diversifying neural progenitor lineages, the underlying molecular mechanisms are poorly understood. Here, using the developing retina as a model system, we addressed the role of nucleosome remodelling in regulating the chronology of cell type production from retinal progenitors. We hypothesized that Chd4 would control the competence windows that define embryonic versus postnatal phases of retinal neurogenesis. In accordance with this hypothesis, we observed a surprisingly straightforward ‘temporal-looking’ shift in neuron production, with *Chd4* cKOs overproducing early-born RGCs and amacrines. Later-born rods were drastically decreased (Fig. 8A, B). While we predicted that this shift in cell composition might arise due to a prolongation in the early competence window, this proved not to be the case. Indeed, scRNA-seq profiling suggested that *Chd4* cKOs were instead slightly accelerated in their temporal state. This latter result matched birthdating experiments showing that rod photoreceptor production wanes precociously. Moreover, we observed that cycling RPCs persisted beyond their normal developmental window. Taken together, these data suggest that Chd4-dependent nucleosome remodelling regulates the temporal transition that terminates the retinal lineage, but does not control earlier competence transitions.

**Fig. 8.**
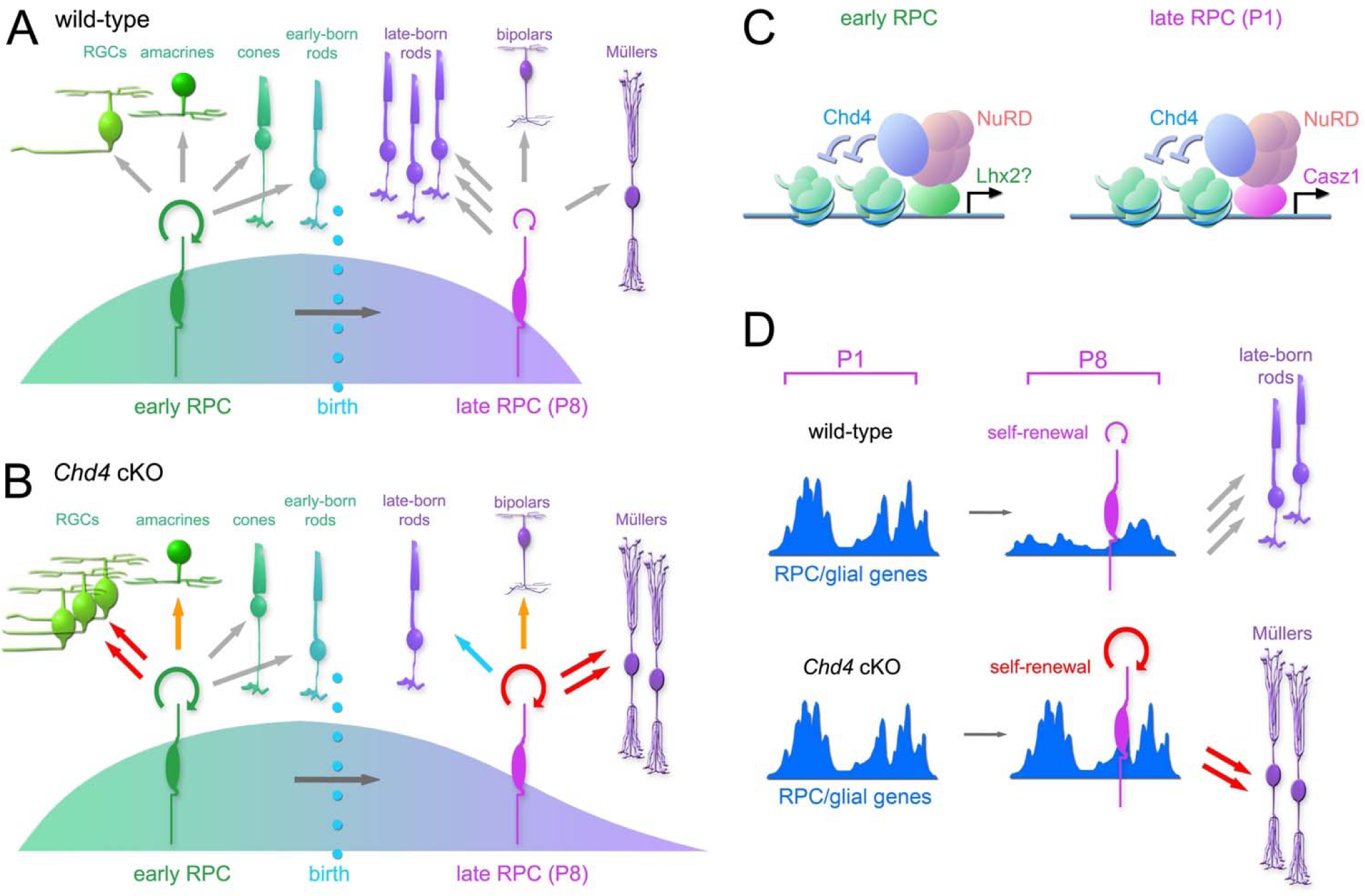
(A, B) Summary of alterations in cell-type production in wild-type versus *Chd4* cKO retinas. (C) Chd4 may associate with different transcription factors at different developmental stages in order to control cell fate specification. (D) Model for role of nucleosome remodelling in temporal progression of RPCs. In the absence of Chd4, expressed genes remain more accessible, reinforcing the progenitor state.

### Molecular control of progenitor competence

Competence determinants were first identified in *Drosophila* neuroblasts [2]. Transcription factor cascades were shown to progressively modify progenitor potential, allowing neuroblasts to produce sequences of different neurons. In vertebrate lineages, epigenetic processes were the first regulators of temporal competence states to be identified [23, 24], but transcription factor cascades that are analogous or even orthologous to *Drosophila* temporal factors were subsequently identified. It remains unclear whether/how transcription factors and heterochromatic determinants converge.

However, multiple temporal transcription factors, including Ikzf1, Casz1, Foxp1, and Lhx2, have been shown to interact with the NuRD complex in various contexts [31–36]. We note that *Chd4* cKOs resemble *Casz1* cKOs, but have a stronger phenotype. Intriguingly, the *Drosophila* orthologue *castor* is similarly required to terminate neuroblast lineages [52], reinforcing the deep evolutionary conservation of this process. In terms of cell type composition, *Chd4* cKOs also resemble *Lhx2* cKOs [17] - perhaps suggesting Lhx2/NuRD cooperativity, which would be consistent with biochemical and functional data from the neocortex [36]. By contrast, the *Ikzf1* mutant underproduces early-born cell types [15, 53], and is in this respect opposite to the *Chd4* cKO, suggesting that Ikzf1 likely acts in a NuRD-independent fashion. Since Ikzf1 regulates early competence, these observations imply that different temporal transitions are governed by different mechanisms – despite the fact that Chd4 is a known co-factor of Ikzf1. Nonetheless, we speculate that dynamically expressed transcription factors may interact with Chd4/NuRD in a stage-dependent fashion to regulate cell fate (Fig. 8C).

In agreement with our results, distinctions between the epigenetic mechanisms that regulate different temporal transitions have been previously described in cortical lineages. Although polycomb was previously shown to regulate multiple competence transitions, including the transition between the production of early versus late-born neurons [25, 54], Tsuboi et al. previously found that NuRD regulated the switch between neurogenesis and gliogenesis but did not regulate earlier temporal transitions [29]. While Nitarska et al. previously showed that Chd4 is specifically required for the production of upper-layer cortical neurons [26] - perhaps suggesting regulation of early-to-late temporal transitions, this phenotype appears to be driven primarily by stage-specific defects in progenitor proliferation. Thus, even though Chd4 has striking effects on the production of early vs. late-born neurons in both the cortex and retina, we conclude that Chd4 -dependent chromatin remodelling does not control the temporal transition that restricts cell type production to embryonic versus postnatal windows.

In cortical lineages, heterochromatic chromatin remodelling complexes such as NuRD and polycomb were previously shown to regulate the switch from neurogenic to gliogenic competence [23, 29]. In agreement with these studies, we observed that Müllers were overproduced in the *Chd4* cKO retina. However, while gliogenesis is analogously upregulated in the *Chd4* cKO retina, there are key differences between the retinal and cortical phenotypes. While glia differentiate precociously in cortical lineages in the absence of NuRD or polycomb, we think that Müller overproduction in the *Chd4* cKO retina occurs indirectly via a failure to exhaust cycling RPCs, rather than through precocious gliogenesis. Indeed, while the production of late-born rods was decreased, bipolar neurons were actually increased in the *Chd4* cKO at P8, albeit that supernumerary bipolars appear to die back by P15. Premature gliogenesis would presumably truncate the normal RPC lineage – leading to a balanced loss in rods and bipolars. Unlike cortical lineages, RPCs do not convert into true glioblasts. Instead, RPC lineages can terminate via Müller differentiation [55], with RPCs perhaps directly converting into Müllers [11, 12]. The observation that cycling RPCs persist in the *Chd4* cKO retina at late stages suggests that this mode of lineage termination becomes favored – probably because terminal divisions that produce rod photoreceptors are undermined. Intriguingly, RPCs in *Nfia/b/x* triple mutants were previously shown to persist beyond the normal termination of neurogenesis, generating supernumerary rods, but failing to generate Müllers [8]. The complementarity of the *Chd4* and *Nfia/b/x* cKO phenotypes might perhaps illustrate two alternative mechanisms that act in parallel to terminate the retinal lineage.

### Role of nucleosome remodelling in retinal progenitors

To understand how Chd4 regulates the genome, we focused on perinatal stages, where birthdating analyses showed that Chd4 was required to promote rod production, and yet RPCs did not yet exhibit alterations in proliferation or apoptosis. Our scRNA-seq results showed that loss of *Chd4* led to a global transcriptomic dysregulation. DEG analysis revealed that loss of *Chd4* affected the expression of various genes involved in cell fate specification. Downregulated genes included the late proneural gene *Ascl1* whereas a number of RPC/glial genes upregulated, likely presaging the later requirement for *Chd4* to promote the exhaustion of RPCs at the end of retinal development.

To directly visualize the nucleosome remodelling activity of Chd4, we performed ATAC-seq on sorted RPCs, revealing that in the absence of *Chd4* there were modest increases in accessibility at thousands of regulatory elements, which correlated with increased transcription at associated genes. Similar functions were previously described in cerebellar granule neurons, where Chd4 depletion led to a widespread increase in genome accessibility [27, 51]. These modest but genome-wide changes in accessibility have also been reported in embryonic stem cells, where NuRD was required to suppress inappropriate gene expression and for the efficient activation of genes required for differentiation [56, 57]. Taken together, our data suggest that without Chd4, the downregulation of RPC gene expression signatures is undermined, leading to a reinforcement of the RPC state (Fig. 8D).

In the developing retina, Chd4 directly occupied approximately 10,000 peaks, similar to what was reported in the cerebellum and neocortex [27, 45]. However, Mbd3 cut&run-seq resulted in only 3,500 peaks, perhaps suggesting NuRD independent functions of Chd4. Recent studies have shown that apart from the NuRD complex, Chd4 can also form a complex with the transcription factor Adnp in order to form the ChAHP complex, which in turn can regulate cortical neurogenesis [45, 58]. The Chd4 phenotype might also arise due to the dysregulation of genome looping. Ctcf was one of the overrepresented footprints in *Chd4* cKO RPCs. Chd4 was previously shown to regulate Ctcf binding and thereby regulate genome architecture [51], and might regulate cohesin through the ChAHP complex, since Adnp and Ctcf have been shown to compete for the same DNA motifs [59]. In the future, it should be interesting to examine *Adnp* cKOs in order to determine how ChAHP regulates retinal development. Future studies will be required to determine whether Chd4 regulates neurogenic competence through NuRD, ChAHP, and/or via interactions with additional co-factors, such as temporal transcription factors.

## Materials and Methods

### Animals

Animal work was conducted according to the guidelines laid out by the Canadian Council of Animal Care under the supervision of the Animal Care and Veterinary Service at the University of Ottawa under the ethical protocols OHRI-2856, OHRI-2867, OHRI-3949, and OHRI-4029. Mice with *Chd4* floxed alleles (*Chd4*^f/f^) were graciously donated by the Katia Georgopoulos laboratory (Harvard, MA, USA) [38], and backcrossed onto the C57BL/6J background (obtained from Jackson Laboratories; RRID: IMSR_JAX:000664). *Chx10-Cre-YFP* transgenic mice (RRID:IMSR_JAX:005105) [39] were generously provided by the Catherine Tsilfidis laboratory. Both males and females were used for all experiments. Genotyping primers are presented in Table S1.

### Histology

For embryonic stages, whole heads were collected and fixed in 4% PFA overnight at 4°C. This was followed by 3 washes in 1X PBS for 5 minutes each followed by immersion in 20% sucrose in PBS at 4°C overnight. The next day they were subjected to 3 washes in 1X PBS for 5 minutes each and submerged in 1:1 solution of OCT:20% sucrose in 1X PBS overnight at 4°C. Subsequently, the whole heads were embedded in the OCT compound and were stored at -80°C. For P0 and P2, a small slit was introduced between the lens and choroid to allow fixative to penetrate the eye. Eyes were then fixed in 4% PFA for 15 minutes. For P8 and P15, lenses were removed to generate eye-cup This was followed by 4% PFA fixation for 2-3 minutes. After fixation, the retinal tissues underwent 3 washes in 1X PBS, and were transferred to 20% sucrose in PBS at 4°C for 2-3 hours. They were finally immersed in the OCT compound for storage at -80°C.

14µm coronal cryosections were collected onto Superfrost Plus slides (Fisher) and processed for immunofluorescence as described previously [21, 35]. Briefly, the tissue sections were washed thrice in 1X PBS followed by antibody incubation at 4°C overnight. Primary antibodies were diluted to 1:100 in blocking buffer (PBS supplemented with 0.4% Triton X-100, 3% w/v BSA, and 1:5000 Hoechst 33342). After washing with 1X PBS, Alexa 555, or 647 –conjugated secondary antibodies were added at a dilution of 1:1000 in blocking buffer and incubated for 2-3 hours at room temperature. Finally, the coverslips were mounted using the Mowiol mounting media (12% w/v Mowiol 4-88, 30% w/v glycerol, 120mM Tris-Cl pH 8.5, 2.5% DABCO) and stored at 4°C until imaged. Antibody information is presented in Table S2.

### Imaging and Cell counting

Images were acquired on an LSM900 confocal microscope (Zeiss), using a 63X objective (Plan-Apochromat 63x/1.40 Oil DIC f/ELYRA) with a 0.5X digital zoom. The images were tiled using Zen software (Zeiss) where required. For each biological replicate, four optical sections (single Z planes) of the peripheral retina were used to generate the cell counts. Manual cell counting was performed using Fiji (ImageJ), wherein each optical section analyzed was cropped to a constant width of 100µm. Images were further processed using Adobe Photoshop CS3 (Adobe) software.

### EdU incorporation

P0 and P1 pups were injected intraperitoneally with 10mM of EdU (Invitrogen #C10640) and eyes were harvested at P2 or P15 as indicated in the figure legends. EdU staining was performed on retinal sections using the Click-iT™ Plus EdU Cell Proliferation Kit for Imaging (Invitrogen #C10640) according to the manufacturer’s protocol. In the instances where EdU was co-stained with other antibodies, the tissues were first processed with immunostaining with primary antibodies before the Click-iT staining reaction.

### Explant Transduction

Retroviral preparation, *ex vivo* retroviral transduction, and clone reconstruction were performed as described previously [21, 35], with minor modifications. *Cre* cDNA was cloned into *pMSCV-EGFP* vectors and used for Cre transduction into *Chd4 f/+* and *Chd4 f/f* retinal explants at P0. The media was changed on a daily basis for 14 days, following which the explants were fixed in 4% PFA and frozen in OCT. Subsequently, the tissues were sectioned and analyzed by IHC. Cell types were annotated using morphology and laminar distribution as described previously [21, 35].

### Statistical analysis

Statistical analysis for image count data was performed using Microsoft Excel and GraphPad Prism version 8 (GraphPad) software. n-values refer to biological replicates and each data point denotes a single biological replicate. For cell-counting, a minimum of 3 biological replicates were quantified, and statistical analysis was performed via one-way ANOVA with Tukey’s multiple comparison test or two-tailed unpaired t-test. We did not perform statistical analyses to predetermine sample sizes. The count data is presented by mean ± standard error of mean (SEM). ∗p < 0.05; ∗∗p < 0.05; ***p < 0.005; ∗∗∗∗p < 0.0005. ns = not significant.

### Western Blot

Western blotting was performed as previously described [35] with some modifications. P0 dissected retinas were homogenized in the RIPA lysis buffer with protease inhibitors (cOmplete, Mini, EDTA-free; 11836170001; Millipore Sigma) and incubated on ice for 15 minutes. Thereafter, they were sonicated on ice using a Cole-Parmer Ultrasonic Homogenizer (RK-04711-45) at 20% amplitude with an 8-sec pulse followed by a 30-sec interval for a total of 3 pulses and centrifuged at max. speed at 4°C for 15 mins. The supernatant was collected, and the protein concentration was quantified using BCA assay. Approximately 40µg of protein lysate was separated on a 6-10% gradient SDS-PAGE gels and semi-dry transferred onto PVDF membranes (Millipore).

### Retinal dissociation

Retinas were dissociated using papain (0.003N NaOH, 100U papain solution (Worthington), 0.4% DNaseI (Millipore Sigma 04716728001), 2mg L-cysteine crystal in 1X PBS) for 8 minutes at 37°C. Subsequently, papain buffer was aspirated and replaced with LO-OVO solution (1X LO-OVO (Bio Basic), 0.4% DNaseI in 1X PBS). Thereafter, the retinal tissue was triturated and centrifuged for 11 minutes at 200g at room temperature. The retinal cells were then re-suspended in 1X PBS.

### ATAC-seq

Two biological replicates were used for each genotype, namely, control (wt, *Chd4*+/+^Cre+^) and mutants (cKO, *Chd4* f/f^Cre+^). P1 retinas from each biological replicate were dissected to make a dissociated suspension as described above. Subsequently, 75,000 YFP+ cells were flow-sorted and used for the ATAC-seq assay as described previously [60, 61] using the Nextera library kit (Illumina).

The analysis of the ATAC-seq dataset was performed using the Galaxy interface [62]. After the initial quality control, the NGS adapters were trimmed using Trimmomatic, and the reads were mapped to the mm9 reference genome using Bowtie2. After merging the alignment files via Samtools merge, peak calling was performed with MACS2. Differential peak detection and analysis were performed using DiffBind and Limma. DiffBind peaks were sorted via k-means clustering with Seqplots and annotated to nearby genes using GREAT. ATAC-seq and cut&run-seq data are available on the GEO database under accession GSE266039.

### Multi-seq

Multi-seq was performed as described previously [43, 61] with some modifications. For this assay, we utilized 3 biological replicates each for control (*Chd4* f/+^Cre+^) and mutant (Chd4 f/f ^Cre+^) samples, for a total of 6 replicates. P1 retinas from each biological replicate were dissociated. Approximately, 250,000 dissociated cells per replicate were then barcoded by incubating with ‘anchor’ and ‘co-anchor’ lipid-modified oligonucleotides graciously provided by the Gartner lab. Barcode oligonucleotides were purchased from Integrated DNA Technologies (see Table S1). Barcodes 1, 3 and 4 were used to tag individual control replicates while barcodes 2, 5 and 6 tagged individual mutant replicates. The respective barcode sequence is shown in the figure panel. Individual replicates were co-incubated with barcode oligonucleotides and pooled into a single tube at a 1:1 ratio. Approximately 20,000 pooled cells were used in a single Chromium™ run (3′ Library & Gel Bead Kit v2, PN-120237, 10X Genomics).

The resulting expression library FASTQs were processed using CellRanger (10X Genomics). The deMULTIplex workflow was used to perform quality control to remove doublets along with cells that lacked the barcodes. Output files were filtered and analyzed using Scanpy version 1.9.190 in Python (Python Core Team n.d.). Genes detected in less than 3 cells were removed from the analysis. Low-quality cells (less than 5,000 genes detected, less than 20,000 reads/counts detected, or more than 0.05% of mitochondrial genes detected) were excluded. The cell types were annotated through an unbiased deep-learning model based on previously published retinal single-cell expression data [8] using scDeepSort version 1.0. Differential gene expression analyses were performed using Scanpy (Wilcoxon signed-rank test), or MAST version 1.24.042. The scRNA-seq data will be available on the GEO database.

## Data availability

ATAC-seq and cut&run-seq data will be made available on the GEO database under accession GSE266039.

The scRNA-seq data will be made available on the GEO database.

## Author contributions

Conceptualization: SS and PM. Investigation: all authors. Data curation and visualization: AF, SS, PM. Writing: original draft: PM and SS. Review and editing: All authors. Supervision and funding: PM.

## Acknowledgements

We thank Michel Cayouette, Fei Chang, and David Picketts for comments on a previous version of this manuscript. We thank Katia Georgopoulos and Toshimi Yoshida for sharing the *Chd4^Flox^*mice. We thank Zev Gartner, Chris McGinnis, David Cook, and Barbara Vanderhyden for generously providing Multi-seq protocols, reagents, and advice. We thank the Michael Dyer and Seth Blackshaw labs for providing datasets upon which this study relied. For scRNA-seq and cut&run-seq experiments, we also thank Katayoun Sheikheleslamy, Caroline Vergette, and Pearl Campbell from the *Stemcore Molecular Biology Core facility*, as well as Chris Porter from the *OHRI Bioinformatics Core Facility*. We thank Chloë van Oostende-Triplet and the *Cell Biology and Image Acquisition Core Facility*, the staff of the uOttawa *Animal Care and Veterinary Service*. We thank members of the Mattar and Picketts labs for their ongoing support and input. This work was generously supported by the Canadian Institutes of Health Research (CIHR) Operating Grants (PJT-166032, PJT-166074), as well as the New Frontiers in Research Fund (NFRFT-2022-00327). SS was generously supported by the David M. Shillito Scholarship in Ophthalmology Research. The project was also supported by an infrastructure grant from the Canada Foundation for Innovation for confocal microscopy (JELF 37688). PM gratefully holds the Gladys and Lorna J. Wood Chair for Research in Vision.

