## Supplemental for "Chd4 remodels chromatin to control retinal cell type specification and lineage termination"

**Supplemental Figures:**

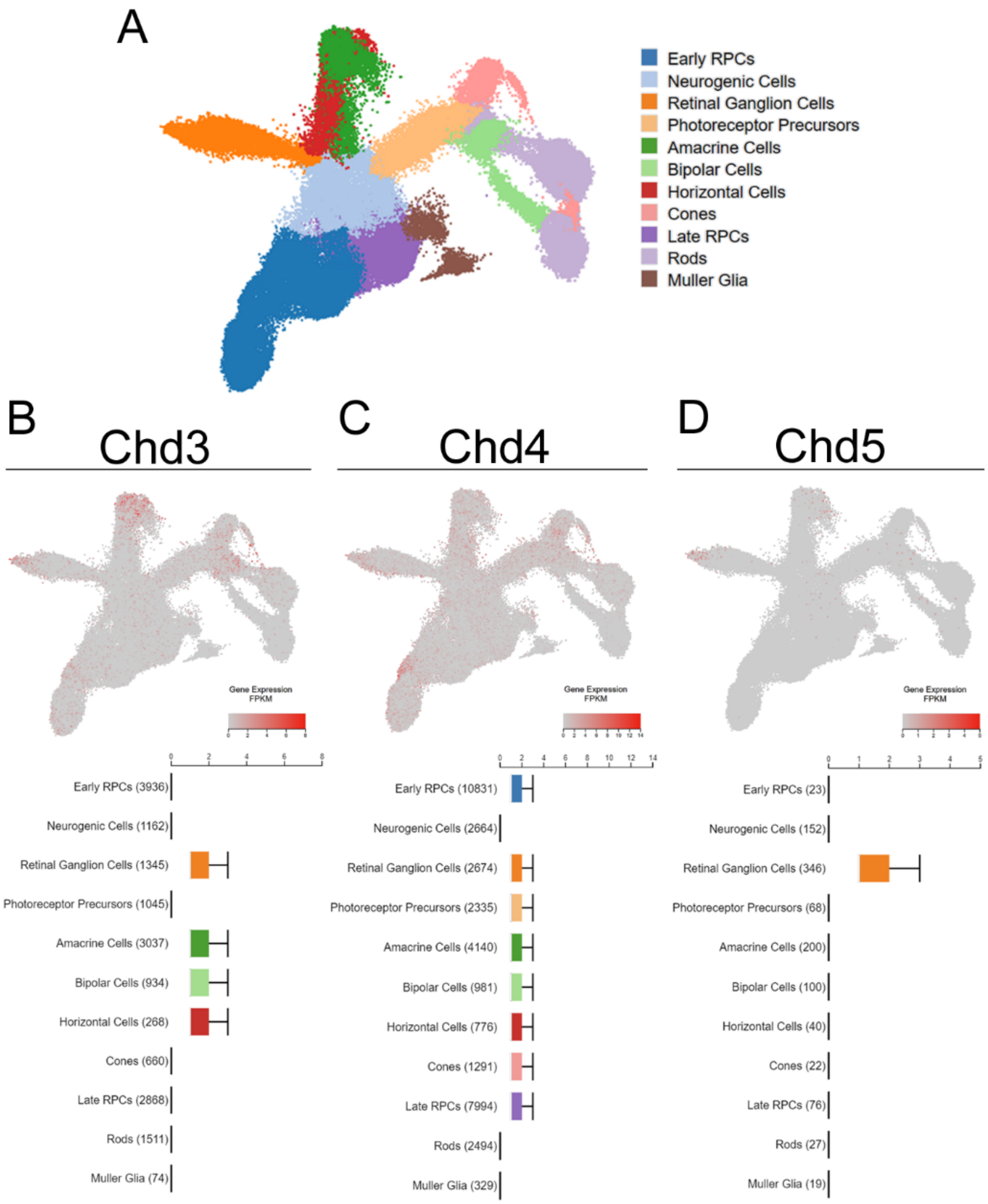


**Fig. S1.** Expression dynamics of *Chd3/4/5* during retinal development. (A) UMAP projection of retinal developmental trajectories from previously published scRNA-seq dataset [8]. (B-D) The expression of individual *Chd* paralogs during retinal development and their respective expression boxplots in different retinal cell-types.

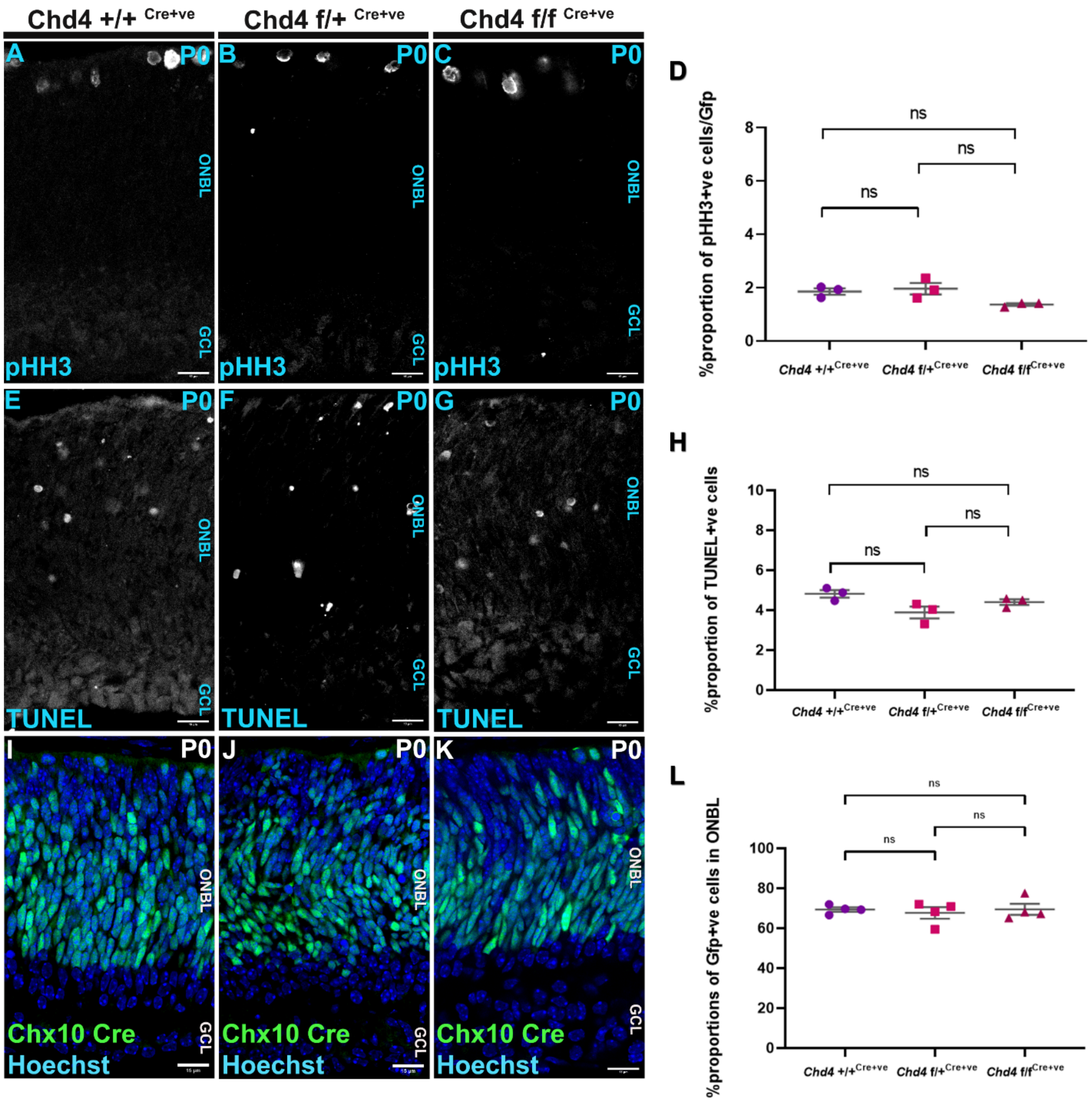


**Fig. S2.** *Chd4* cKO does not affect proliferation or viability at P0. (A-C) Wt, chet, and cKO retinal sections were stained with the mitotic marker phospho-histone H3. (D) Percentage of pHH3+ cells at P0. (E-H) TUNEL assay to quantify apoptotic cells. (I-K) YFP staining to mark Chx10+ progenitors. (L) Using YFP as a proxy for RPCs, the progenitor pool as quantified between the different genotypes. All data are presented as mean ±SEM. ns: not significant by one-way ANOVA with Tukey's multiple comparisons test. Scale bar = 10 microns.

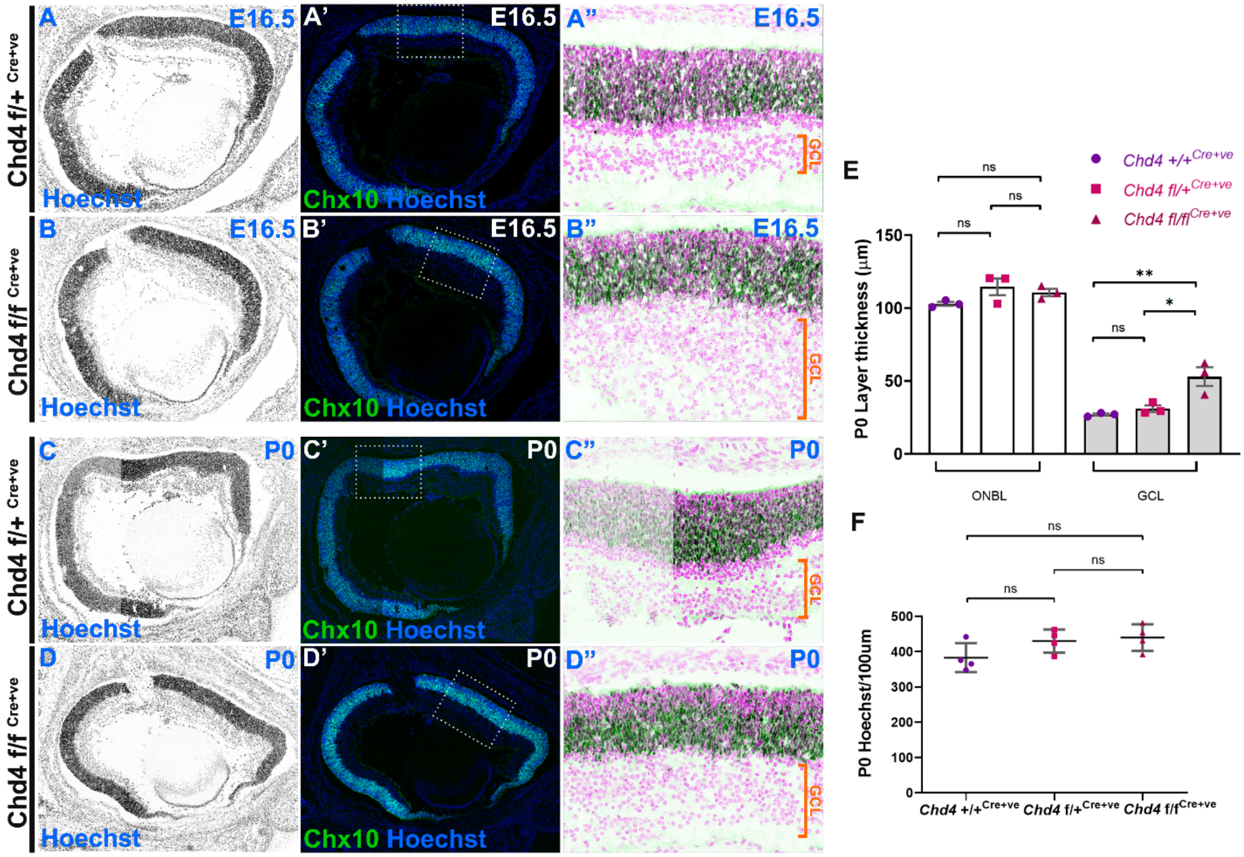


**Fig. S3.** *Chd4* cKO leads to embryonic expansion of the ganglion cell layer and lamination defects. (A-B″) E16.5 chet and cKO whole eye sections were stained with the Hoechst and YFP. (C-D″) P0 chet and cKO whole eye sections were stained with Hoechst and YFP. (E) Quantification of individual layer thickness between the different genotypes. (F) Total Hoechst was quantified and compared across the three genotypes. All data are presented as mean ±SEM. p-value *p<0.05, **p<0.005, ns: not significant by one-way ANOVA with Tukey's multiple comparisons test.

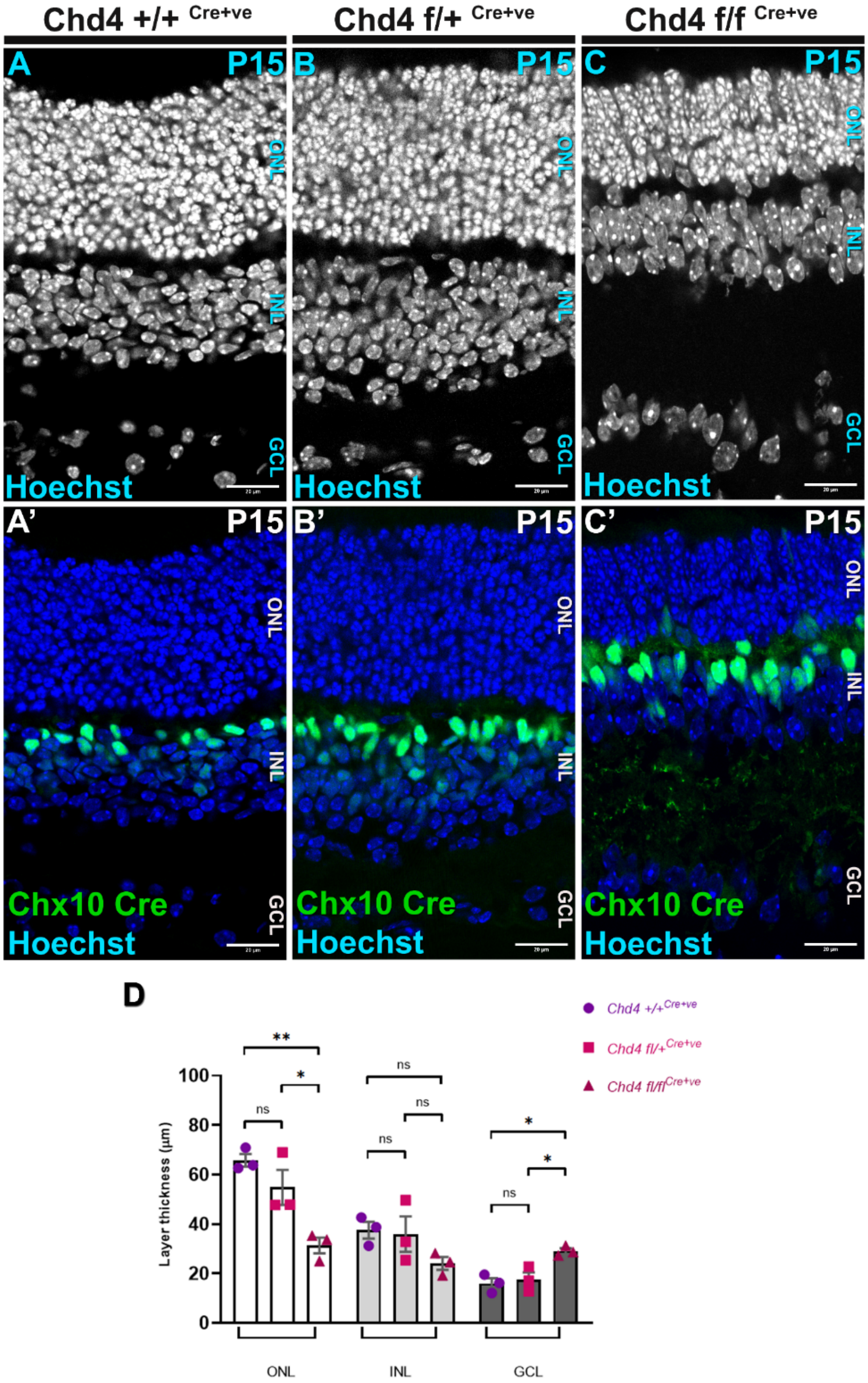


**Fig. S4.** Effect of *Chd4* cKO on P15 retinal histology. (A-C') Wt, chet and cKO retinal sections were stained with Hoechst to mark the cell bodies in the different retinal layers. (D) Quantification of individual layer thickness between the different genotypes. All data are presented as mean ±SEM. p-value *p<0.05, **p<0.005, ns=not significant by one-way ANOVA with Tukey's multiple comparisons test. Scale bar = 10 microns.

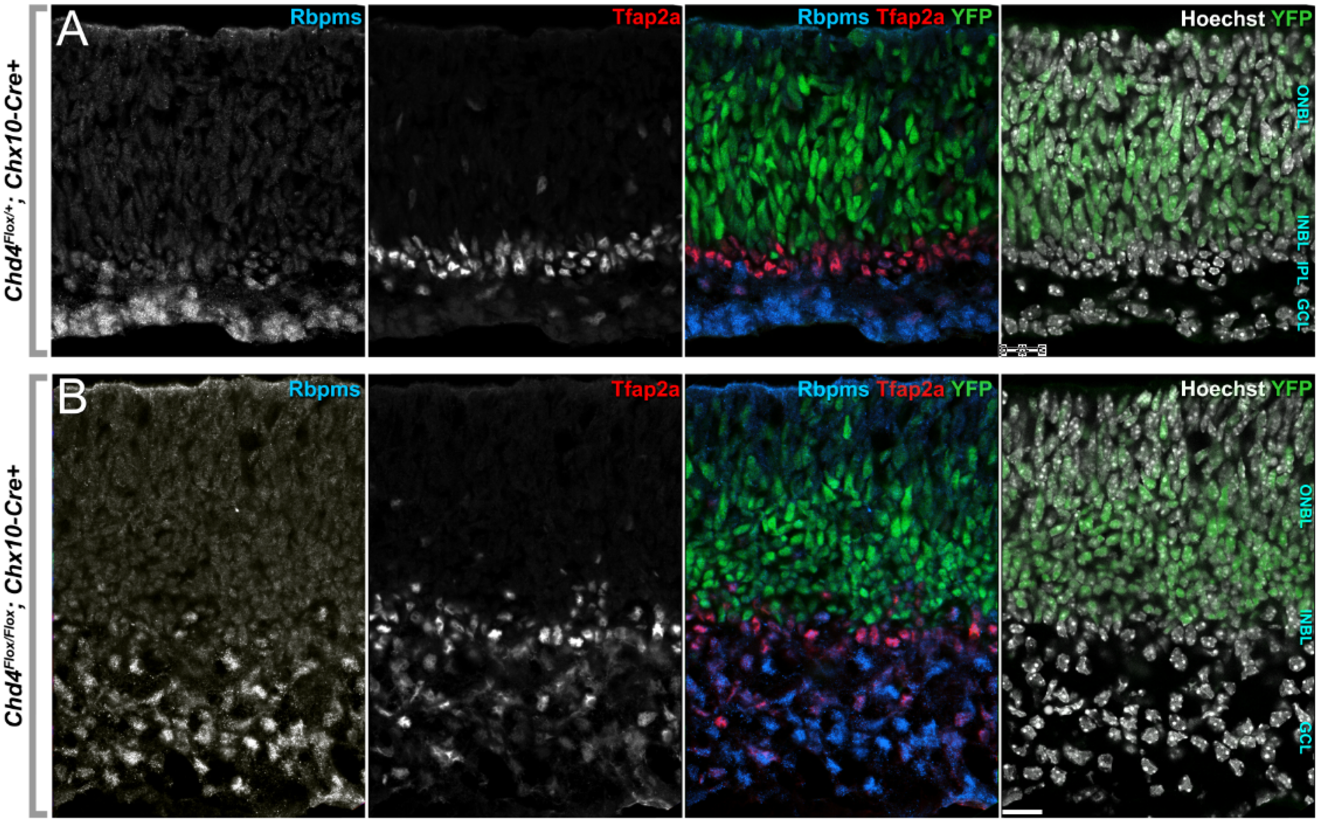


**Fig. S5.** Expansion of ganglion and amacrine cells in the *Chd4* cKO at P0. (A, B) chet (A) and cKO (B) retinas were stained with the ganglion cell marker Rbpms, or the amacrine marker Tfap2a.

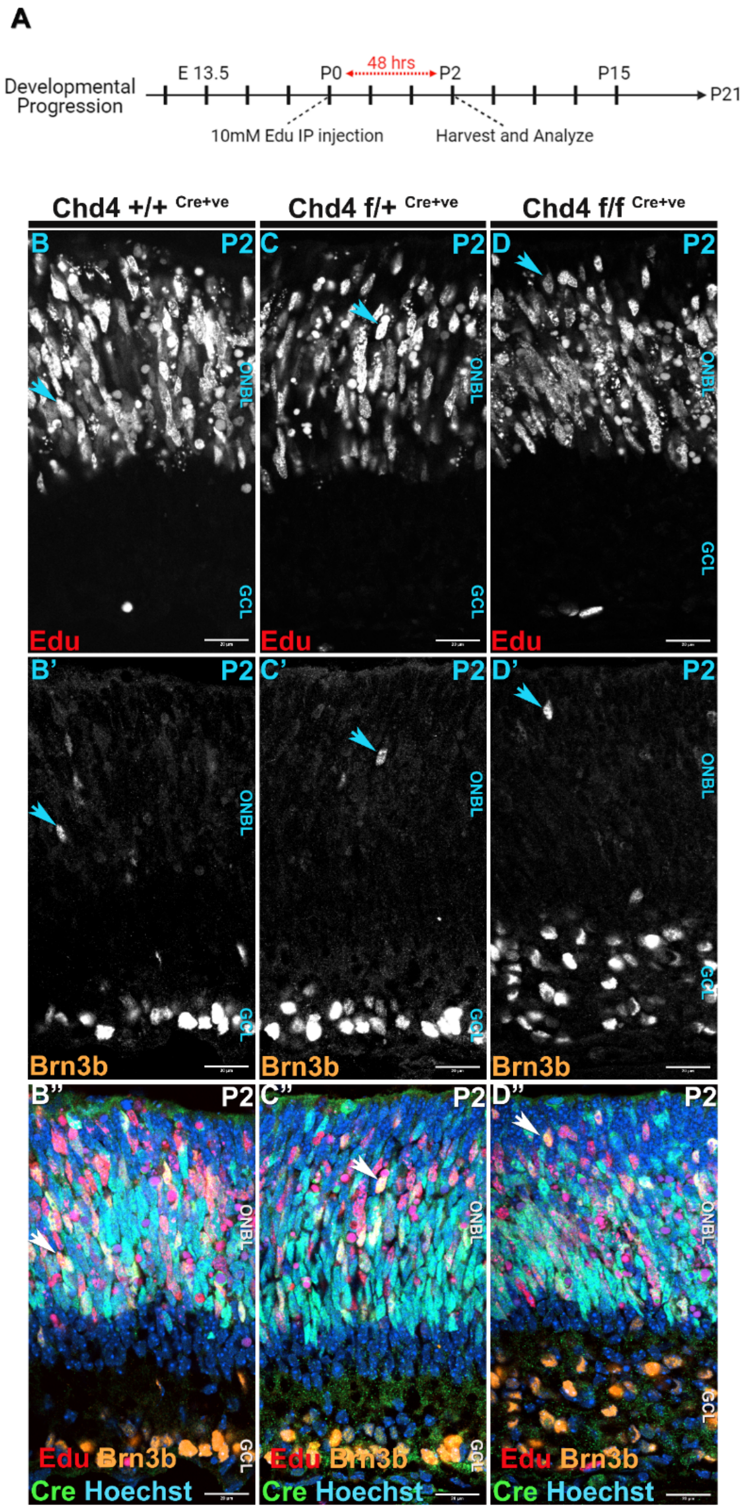


**Fig. S6.** RGCs are overproduced within their normal birth window in *Chd4* cKO retinas. (A) Schematic of 48-hours Edu birthdating assay. (B-D′′) Wt, chet, and cKO retinas were stained with EdU and and Brn3b. Arrows indicate cells that are double positive for EdU and Brn3b. Scale bar = 10 microns.

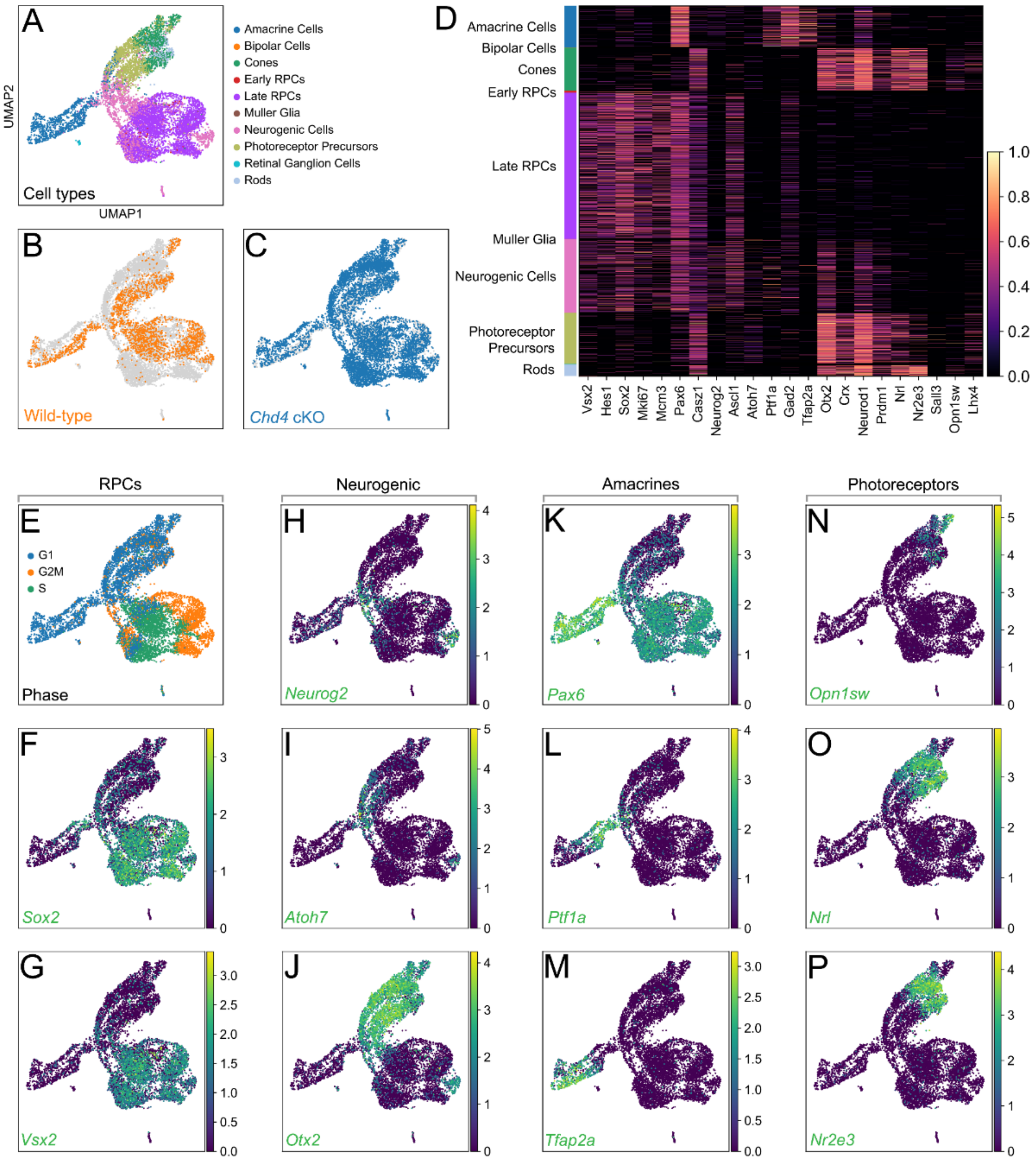


**Fig. S7.** Annotation of UMAP clusters using marker gene expression. (A) Cell-type annotation using label transfer from a published retinal scRNA-seq atlas [8]. (B-C) Segregating the UMAP projection based on cell genotype. (D) Heat map of the expression of marker genes used to annotate the different cell type clusters present in the P1 scRNA-seq dataset. (E-P) UMAP projections of marker gene expression in control and *Chd4* cKO replicates. (E-G) RPC markers are based on the cell-cycle phase (E), along with the expression of progenitor-specific markers *Sox2* (F) and *Vsx2* (G). (H-J) Markers of neurogenic cells determined by the expression of *Neurog2* (H), *Atoh7* (I), and *Otx2* (J). (K-M) Amacrine cell type markers *Pax6* (K), *Ptf1a* (L), and *Tfap2a* (M). (N-P) Photoreceptor markers *Opn1sw* (N), *Nrl* (O), and Nr2e3 (P).

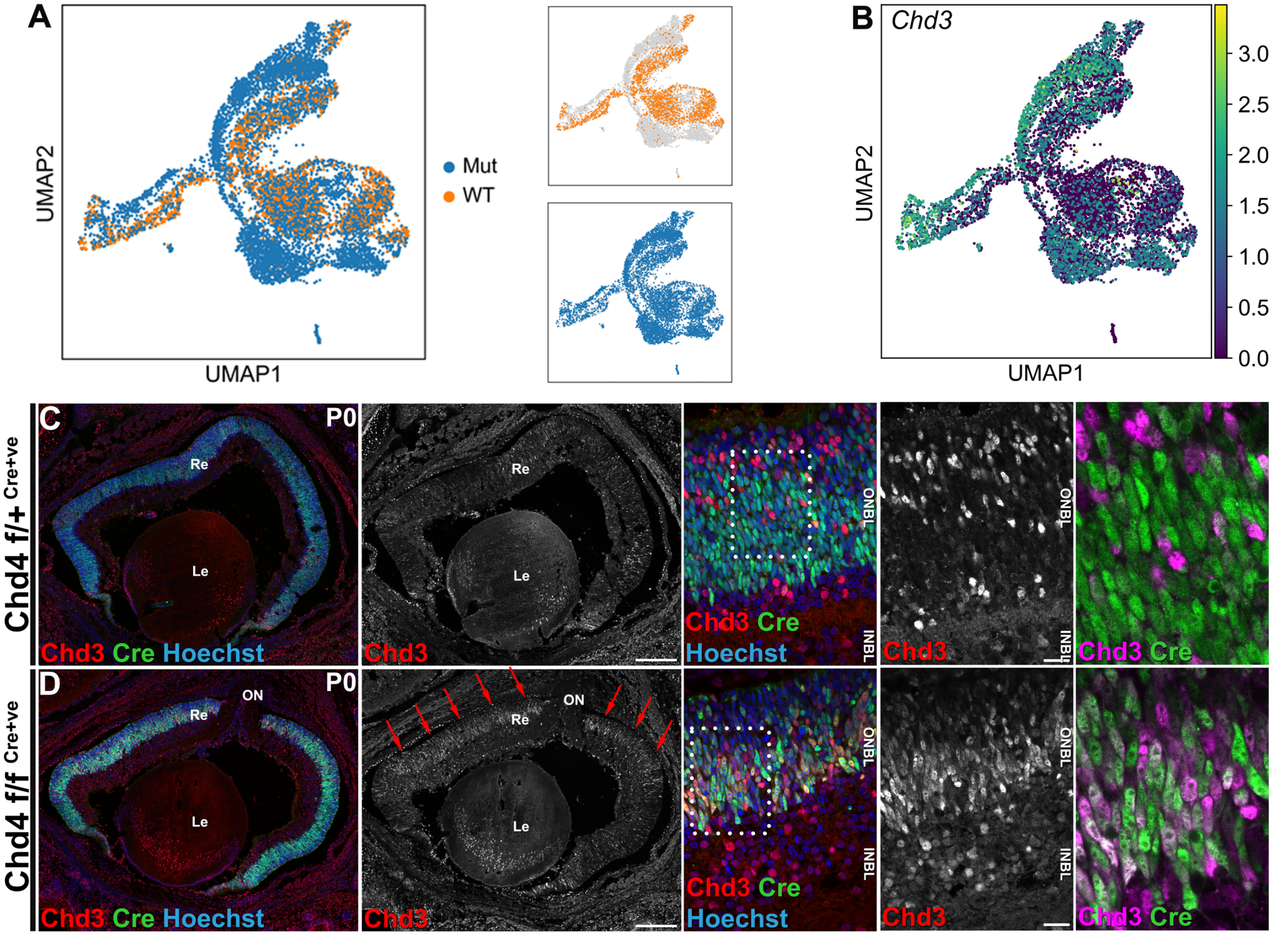


**Fig. S8.** Validation of the upregulation in Chd3 expression in the absence of *Chd4*. (A) UMAP representation of clusters segregated by genotype. (B) UMAP projection of *Chd3* expression in control and cKO samples. (C-D) IHC staining of Chd3 in chet and cKO P1 retinal sections.

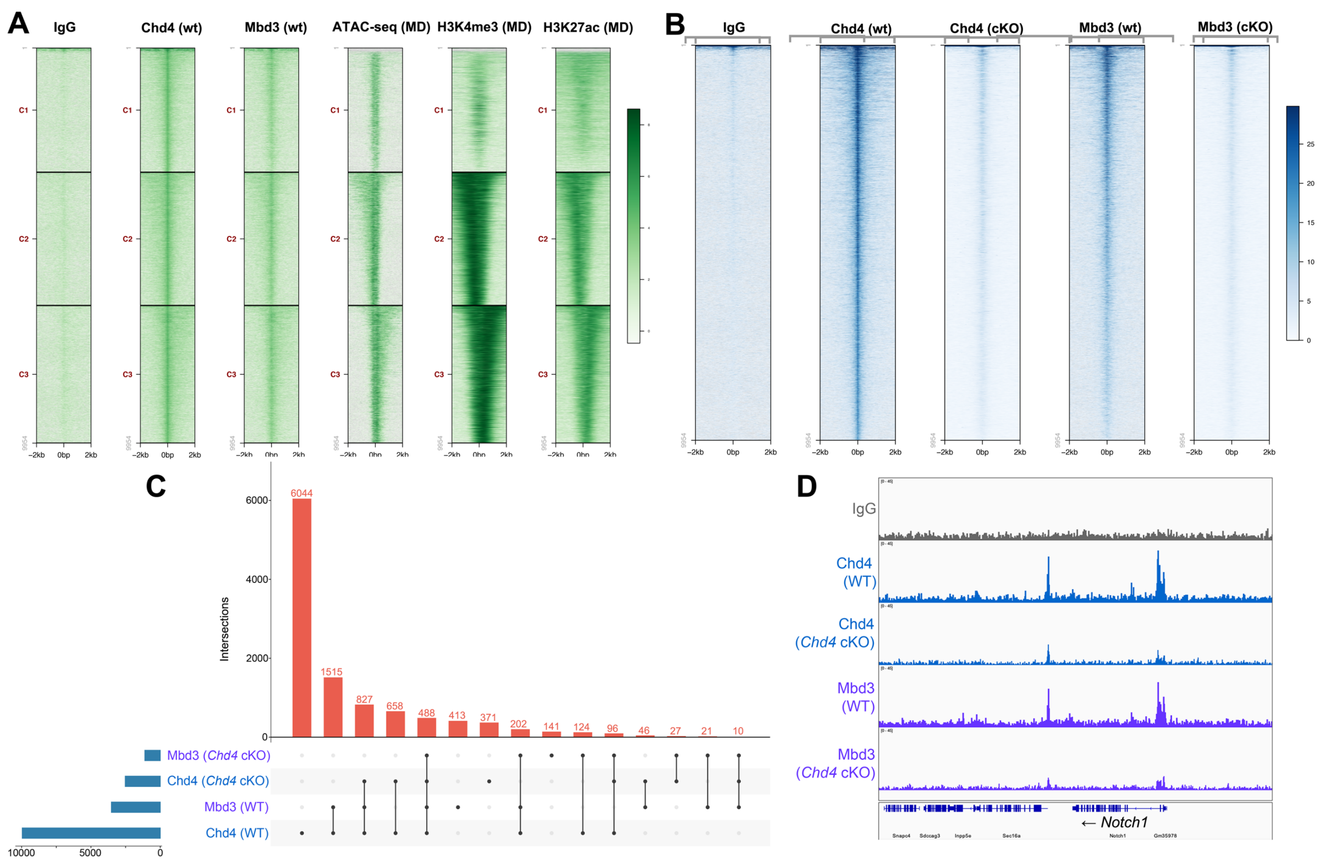


**Fig. S9.** Genomic occupancy of Chd4. (A) Heat map of Cut&Run-seq peaks of IgG, Chd4 and Mbd3 from wild-type and *Chd4* cKO P1 retinas centered on Chd4 wild-type peaks. (B) Comparing the wild-type cut&run-seq dataset with previously published ChIP-seq data[59] on age-matched retinas to determine the genomic occupancy of Chd4 and Mbd3. (C) Upset plot of cut&run-seq peak intersections. (D) IGV track of cut&run-seq peaks on the *Notch1* locus.

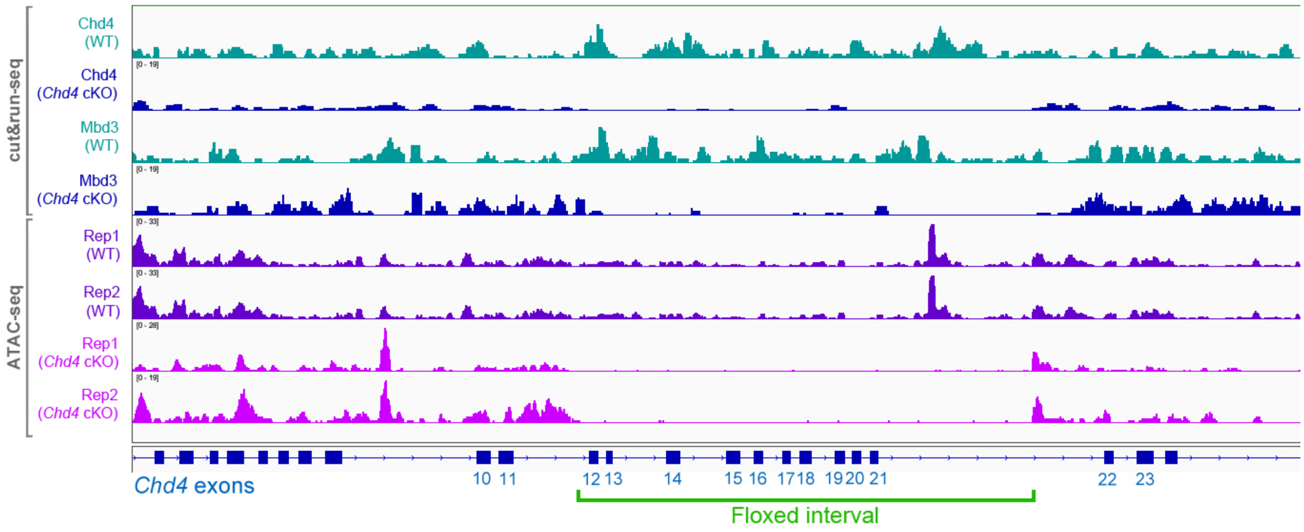


**Fig. S10.** Excision of the loxp flanked region in *Chd4* cKO genomic data.

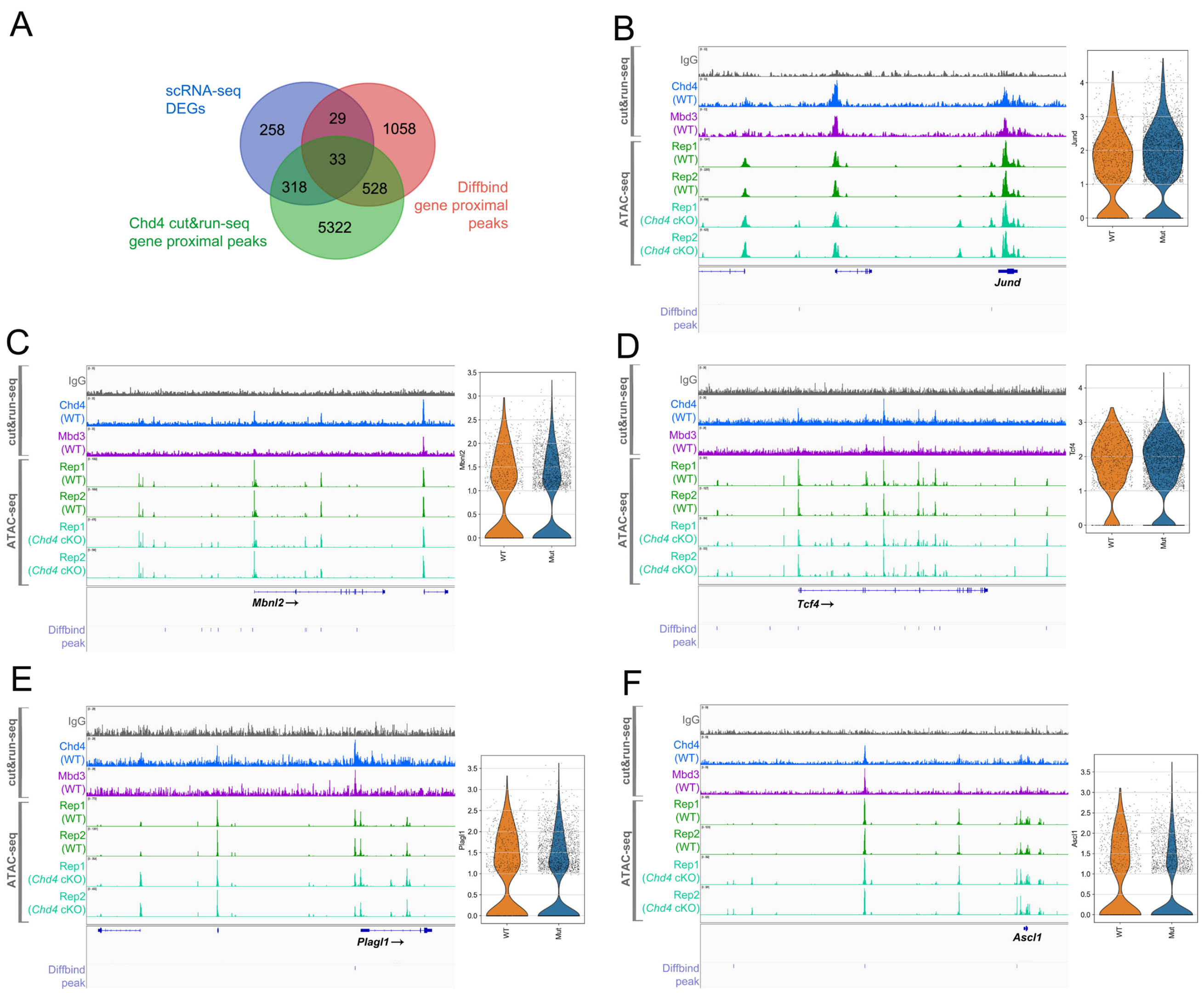


**Fig. S11.** (A) Integration of DEGs from the scRNA-seq analysis with genes associated with gene-proximal DARs and Chd4 Cut&Run-seq gene-proximal peaks. (B) Cut&Run-seq and ATAC-seq tracks for selected DEGs that are directly occupied by Chd4. Diffbind peaks indicate DARs from control vs. *Chd4* cKO ATAC-seq data. Violin plots display scRNA-seq expression data from late RPCs.

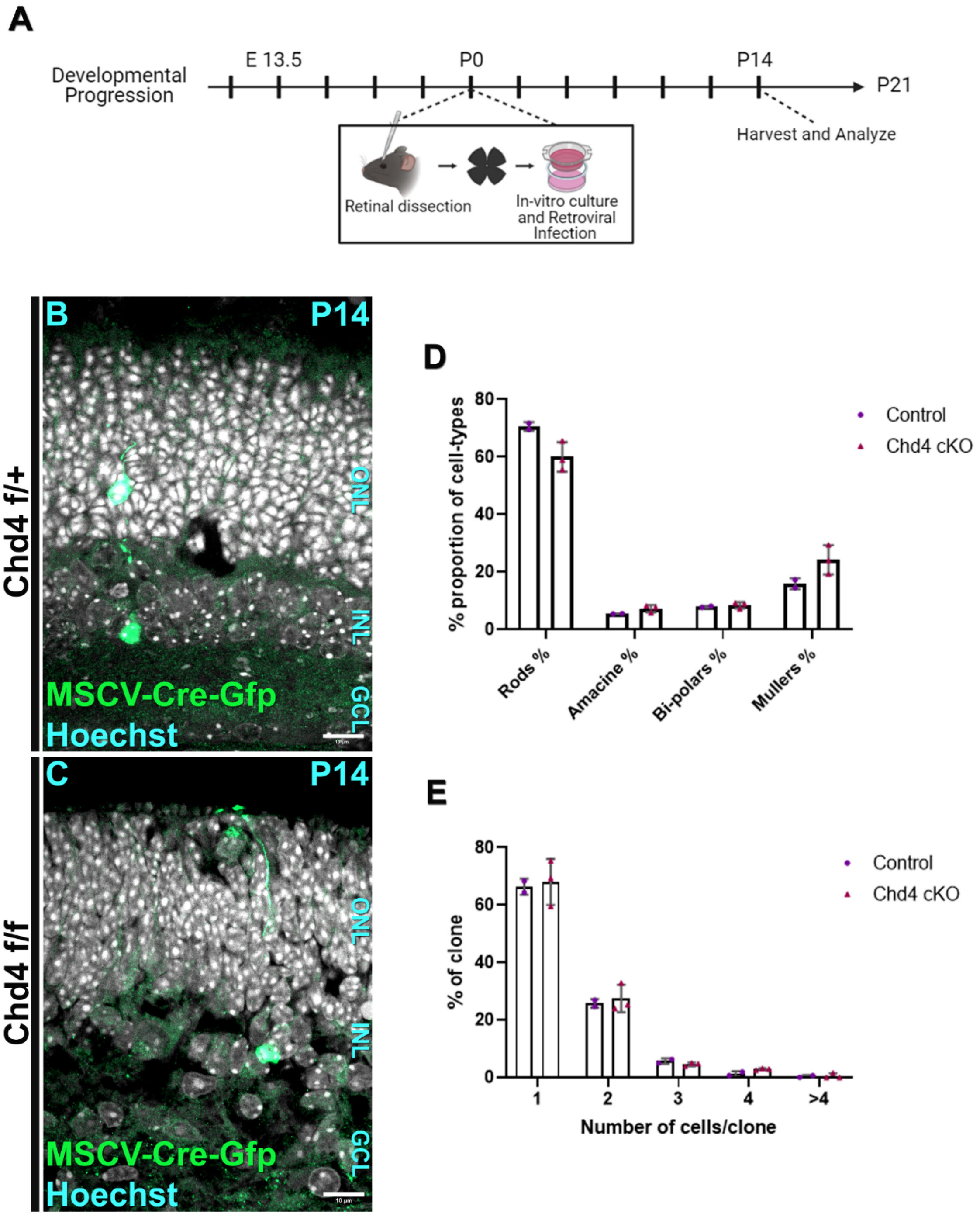


**Fig. S12.** Lineage tracing of *Chd4* cKO RPCs. (A) Experimental plan of the 14-day lineage tracing workflow. (B-C) IHC staining of Gfp and Hoechst delineating RPC-derived clones in *Chd4 f/+* (B) versus *Chd4 f/f* (C). (D) Quantifying the different cell-types generated in control (n=2) and *Chd4* cKO (n=3) clones. (E) Quantifying the cell number per clone in control (n=2) vs *Chd4* cKO (n=3). Scale bar = 5 microns.

**Table S1:** Oligonucleotide sequences. Multi-seq barcodes are highlighted in green.

| Name | Purpose | Sequence |
|---|---|---|
| Mi-2β+F S | genotyping | 5′-CTCCAAGAAGAAGACGGCAGATCT-3′ |
| Mi-2 INR A | genotyping | 5′-GTCCTTCCAAGAAGAGCAAG-3′ |
| CRE-F | genotyping | 5′-AGGTGTAGAGAAGGCACTTAGC-3′ |
| CRE-R | genotyping | 5′-CTAATCGCCATCTTCCAGCAGG-3′ |
| Multiseq-1 | barcoding | 5′-CCTTGGCACCCGAGAATTCCAGGAGAAGAAAAAAAAAAAAAAAAAAAAAAAAAAAAAAA-3′ |
| Multiseq-2 | barcoding | 5′-CCTTGGCACCCGAGAATTCCACCACAATGAAAAAAAAAAAAAAAAAAAAAAAAAAAAAA-3′ |
| Multiseq-3 | barcoding | 5′-CCTTGGCACCCGAGAATTCCATGAGACCTAAAAAAAAAAAAAAAAAAAAAAAAAAAAAA-3′ |
| Multiseq-4 | barcoding | 5′-CCTTGGCACCCGAGAATTCCAGCACACGCAAAAAAAAAAAAAAAAAAAAAAAAAAAAAA-3′ |
| Multiseq-5 | barcoding | 5′-CCTTGGCACCCGAGAATTCCAAGAGAGAGAAAAAAAAAAAAAAAAAAAAAAAAAAAAAA-3′ |
| Multiseq-6 | barcoding | 5′-CCTTGGCACCCGAGAATTCCATCACAGCAAAAAAAAAAAAAAAAAAAAAAAAAAAAAAA-3′ |

**Table S2:** Antibodies and dilutions.

| Antigen | Species | Dilution | Supplier | Catalog | RRID | Application |
|---|---|---|---|---|---|---|
| Brn3a | Mouse | 1:100 | Millipore Sigma | MAB1585MI | AB_94166 | IHC |
| Brn3b | Goat | 1:200 | Santa Cruz | sc-6026 | AB_673441 | IHC |
| Chd3 | Rabbit | 1:200 | Fortis | A301-220A-T | AB_890570 | IHC |
| Chd4 | Rat | 1:200 | Biolegend | 942302 | AB_2888898 | IHC |
| Chd4 | Rabbit | 1:1000<br>1:100 | Abcam | ab72418 | AB_1268107 | Western, cut&run |
| Cleaved Caspase-3 (Asp175) | Rabbit | 1:500 | Cell Signaling | 9579 | AB_10897512 | IHC |
| Cone arrestin | Rabbit | 1:200 | Millipore Sigma | AB15282 | AB_1163387 | IHC |
| GFP | Mouse | 1:100 | DSHB | GFP-G1 | AB_2619561 | IHC |
| Ki67 | Rabbit | 1:100 | Millipore Sigma | SAB5500134 | AB_2892217 | IHC |
| Lhx2 | Rabbit | 1:200 | Millipore Sigma | ABE1402 | AB_2722523 | IHC |
| Otx2 | Goat | 1:500 | R&D Systems | AF1979 | AB_2157172 | IHC |
| Pax6 | Rabbit | 1:500 | Novus | NBP2-19711 | AB_3264565 | IHC |
| Pax6 | Rabbit | 1:500 | Proteintech | 12323-1-AP | AB_2159695 | IHC |
| pHH3 | Rabbit | 1:500 | Cell Signaling | 9701S | AB_331535 | IHC |
| Rbpms | Guinea pig | 1:200 | Millipore Sigma | ABN1376 | AB_2687403 | IHC |
| Rxrγ | Mouse | 1:100 | Santa Cruz | sc-365252 | AB_10850062 | IHC |
| Sox2 | Goat | 1:500 | R&D Systems | AF2018-SP | AB_355110 | IHC |
| Tfap2a | Mouse | 1:500 | DSHB | 3B5 | AB_2313948 | IHC |
| anti-guinea pig Alexa Fluor™ 555 | Goat | 1:1000 | Rockland | 606-142-129 | AB_1961616 | 2º (IHC) |
| anti-goat Alexa Fluor™ 488 | Donkey | 1:1000 | Jackson | 705-547-003 | AB_2340431 | 2º (IHC) |
| anti-goat DyLight™ 650 | Donkey | 1:1000 | Novus | NBP1-75604 | AB_11018252 | 2º (IHC) |
| anti-mouse Alexa Fluor™ 555 | Donkey | 1:1000 | Invitrogen | A-31570 | AB_2536180 | 2º (IHC) |
| anti-mouse Alexa Fluor™ 647 | Donkey | 1:1000 | Invitrogen | A-31571 | AB_162542 | 2º (IHC) |
| anti-rabbit Alexa Fluor™ 488 | Donkey | 1:1000 | Jackson | 111-545-003 | AB_2338046 | 2º (IHC) |
| anti-rabbit Alexa Fluor™ 647 | Donkey | 1:1000 | Jackson | 711-607-003 | AB_2340626 | 2º (IHC) |
| anti-rat DyLight™ 550 | Donkey | 1:1000 | Invitrogen | SA510027 | AB_2556607 | 2º (IHC) |
| anti-Rabbit HRP | Donkey | 1:10 000 | GE Healthcare | NA934 | AB_772206 | 2º (Western) |
